# Tankyrase inhibition sensitizes melanoma to PD-1 immune checkpoint blockade in syngeneic mouse models

**DOI:** 10.1101/526343

**Authors:** Jo Waaler, Line Mygland, Anders Tveita, Martin Frank Strand, Nina Therese Solberg, Petter Angell Olsen, Kaja Lund, Shoshy Alam Brinch, Max Lycke, Elisabeth Dybing, Vegard Nygaard, Sigurd Læines Bøe, Karen-Marie Heinz, Eivind Hovig, Clara Hammarström, Alexandre Corthay, Stefan Krauss

**Affiliations:** Department of Immunology and Transfusion Medicine, Oslo University Hospital, Oslo, Norway; Hybrid Technology Hub - Centre of Excellence, Institute of Basic Medical Sciences, University of Oslo, Oslo, Norway; Department of Health Sciences, Kristiania University College, Oslo, Norway; Department of Tumor Biology, Institute for Cancer Research, Oslo University Hospital, Oslo, Norway; Department of Medical Biochemistry, Oslo University Hospital, Radiumhospitalet, Oslo, Norway; Center of Bioinformatics, Department of Informatics, University of Oslo, Oslo, Norway; Department of Pathology, Oslo University Hospital, Oslo, Norway

## Abstract

The development of immune checkpoint inhibitors represents a major breakthrough in cancer therapy. Nevertheless, a substantial number of patients fail to respond to checkpoint pathway blockade^1,2^. β-catenin is the key transcriptional regulator of WNT/β-catenin signaling^3^. Evidence for β-catenin-mediated immune evasion is found in 13% of all cancers^4^, 42% of primary cutaneous melanoma^5^ and a mouse melanoma model^6^. Currently, there are no therapeutic strategies available for targeting WNT/β-catenin signaling to counteract checkpoint inhibitor resistance in melanoma. Here we report that a specific small-molecule tankyrase inhibitor, G007-LK^7–9^, attenuates WNT/β-catenin and YAP signaling pathways in the syngeneic murine B16-F10 melanoma model enabling sensitivity to anti-PD-1 immune checkpoint therapy. RNA sequencing of 18 tankyrase inhibitor-treated human melanoma cell lines and B16-F10 cells revealed a transcriptional response profile for a subpopulation. This cell line sub-group displayed elevated baseline YAP signaling activity and was susceptible to reduce melanocyte inducing transcription factor (*MITF*) expression upon tankyrase inhibition.

Checkpoint inhibitor treatment, including blockade of the PD-1 receptor, has shown limited efficacy against B16-F10 tumors despite strong expression of the ligand PD-L1 on the tumor cells^10^, a feature attributed to low infiltration by effector CD8^+^ T cells^11,12^. G007-LK^7–9^ is a potent and specific tankyrase inhibitor that obstructs WNT/β-catenin and YAP signaling through stabilization of the AXIN-based degradosome and AMOT proteins, respectively^8,13–17^. The efficacy of G007-LK treatment against WNT/β-catenin and YAP signaling in wild-type B16-F10 cells was explored *in vitro* and *in vivo*. Limited impact on proliferation was observed in cultured B16-F10 cells upon G007-LK treatment (Supplementary Fig. 1a). G007-LK-treated B16-F10 cells displayed a shift in TNKS1/2 gel-migration *in vitro* and formation of cytoplasmic TNKS1/2-containing puncta reflecting degradosome accumulation^8,15,18^ (Supplementary Fig. 1b,c). In addition, G007-LK reduced the level of β-catenin protein and decreased transcription of WNT/β-catenin and YAP signaling target genes as well as luciferase-based reporter activities (Fig. 1a and Supplementary Fig. 1c-h). Next, B6N-Tyr^c-Brd^/BrdCrlCrl mice (C57BL/6N) challenged with B16-F10 tumors were treated with G007-LK. This treatment significantly decreased β-catenin protein levels in the tumor and attenuated transcription of WNT/β-catenin and YAP signaling target genes (Fig. 1a,b and Supplementary Fig. 2). The results show that tankyrase inhibitor treatment can attenuate WNT/β-catenin and YAP signaling in B16-F10 cells *in vitro* and *in vivo*.

**Fig. 1.**
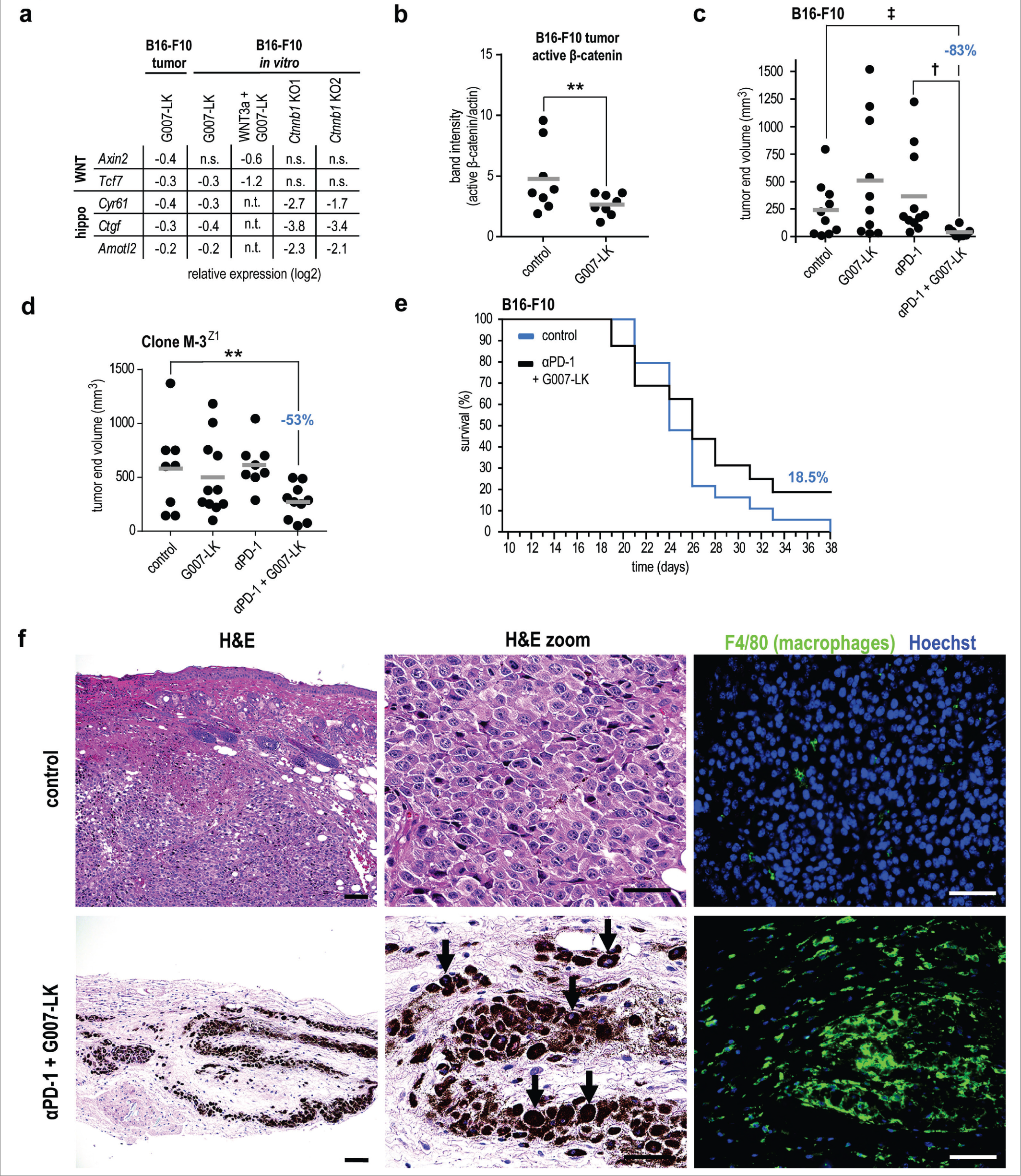
Tankyrase inhibition attenuates WNT/β-catenin and YAP signaling activities in B16-F10 *in vitro* and *in vivo* and confers anti-tumor efficacy in combination with PD-1 inhibitor treatment in mouse models for melanoma. **a**, Relative expression values (log2) from real-time RT-qPCR analyses of WNT/β-catenin (*Axin2* and *Tcf7*) and YAP (*Cyr61*, *Ctgf* and *Amotl2*) signaling target genes. From left to right: G007-LK-treated B16-F10 tumors (subcutaneous [s.c], relative to control), cultured B16-F10 cells treated with G007-LK (1 μM, relative to control), G007-LK/WNT3a (relative to WNT3a-stimulated control) and B16-F10^Ctnnb1KO^ cells (KO1 and KO2, relative to wild-type control). Mean values from 2 or more independent experiments are shown, n.s. = not significant and n.t. = not tested. **b**, Quantification of immunoblotted active β-catenin (non-phosphorylated in Ser33/37/Thr41) from whole s.c. B16-F10 tumor lysates upon 4 days of treatment with G007-LK compared to control (both groups, *n* = 8). One-tailed t-test is indicated by ** (*P* < 0.05). For **b-d**: Mean values are indicated by grey lines. Absence of depicted statistical comparisons indicates lack of statistical significance. **c**, B16-F10 tumor (s.c.) end volume reduction upon anti-PD-1/G007-LK treatment (−83% when compared to control) in mice treated from day 10 through day 21. Control diet (*n* = 10), G007-LK diet (*n* = 10), anti-PD-1 (*n* = 11) and anti-PD-1/G007-LK (*n* = 8). Mann-Whitney rank sum tests are indicated by ^‡^ (*P* < 0.05) and ^†^ (*P* < 0.01). **d**, Clone-M3^Z1^ tumor (s.c.) end volume reduction upon anti-PD-1/G007-LK treatment (−53% when compared to control) in mice treated from day 8 until day 18. Control diet (*n* = 8), G007-LK diet (*n* = 8), anti-PD-1 (*n* = 11) and anti-PD-1/G007-LK (*n* = 10). Two-tailed t-test is indicated by ** (*P* < 0.05). **e**, Kaplan-Meier plot showing survival of B16-F10-recipient (s.c.) mice treated with control diet (*n* = 19, blue) or anti-PD-1/G007-LK (*n* = 16, black) from day 10 through day 38. Difference between the two graphs: One-tailed log rank (Mantel-Cox) test, *P* = 0.087, hazard ratio = 1.75, 95% CI: 0.78-3.93. **f**, Representative pictures from multiple sections of H&E (left and mid panels) and F4/80 and Hoechst-stained (in green and blue, respectively, right panels) tumors from control (*n* = 5, upper panels) and tumor implant sites of anti-PD-1/G007-LK-treated surviving mice (*n* = 2, lower panels). Macrophages laden with melanin are highlighted with arrows. Scale bars: H&E = 100 μm (original magnification ×100) and H&E higher magnification and F4/80 Hoechst = 50 μm (original magnification × 400).

To test whether tankyrase inhibition can counteract resistance to PD-1 immune checkpoint blockade, subcutaneous B16-F10 tumors were established in C57BL/6N mice. Monotherapy with G007-LK or anti-PD-1 resulted in no tumor size reduction, however, combined anti-PD-1/G007-LK treatment resulted in a significant reduction in tumor volume and weight (Fig. 1c and Supplementary Fig. 3). In addition, a significant reduction in tumor volume was also observed upon combined anti-PD-1/G007-LK treatment of murine Clone M-3^Z1^ melanoma in immunocompetent DBA/2N mice (Fig. 1d and Supplementary Fig. 4). No signs of toxicity, intestinal injury (Supplementary Fig. 3g) or body weight changes were observed in any of the mouse experiments (Supplementary Fig. 3f and 4b). To examine longer-term efficacy of combined anti-PD-1/G007-LK treatment, B16-F10-bearing C57BL/6N mice were followed until the entire control group reached the endpoint criterion. In the 3 surviving anti-PD-1/G007-LK-treated mice (18.5%)(Fig. 1e and Supplementary Fig. 5), histopathological inspection of immunostained tumor sections detected no viable tumor cells. Instead, the tumor implant site was infiltrated by macrophages with ingested melanin, presumably derived from B16-F10 cells (Fig. 1f). In summary, the tested murine melanoma models are resistant to single-agent anti-PD-1 or G007-LK treatment, however, a synergistic anti-tumor effect and eradication of a subset of the tumors was observed upon combined anti-PD-1/G007-LK treatment.

We next pursued the mechanistic basis for the observed synergy of anti-PD-1 and G007-LK treatment. To evaluate β-catenin-mediated immune evasion in the B16-F10 syngeneic mouse melanoma model, β-catenin was knocked out in B16-F10 cells (B16-F10^*Ctnnb1*KO^, Supplementary Fig. 6) and the cell line was used to establish subcutaneous tumors in C57BL/6N mice. Compared to vehicle control, anti-PD-1-treated mice displayed significant reduction in tumor sizes, indicating loss of anti-PD-1 resistance in β-catenin-deficient tumors (Fig. 2a and Supplementary Fig. 7). The result suggests that the synergistic anti-PD-1/G007-LK treatment effect (Fig. 1c) can be attributed to G007-LK-induced reduction of β-catenin levels in β-catenin wild-type B16-F10 tumors (Fig. 1b).

**Fig. 2.**
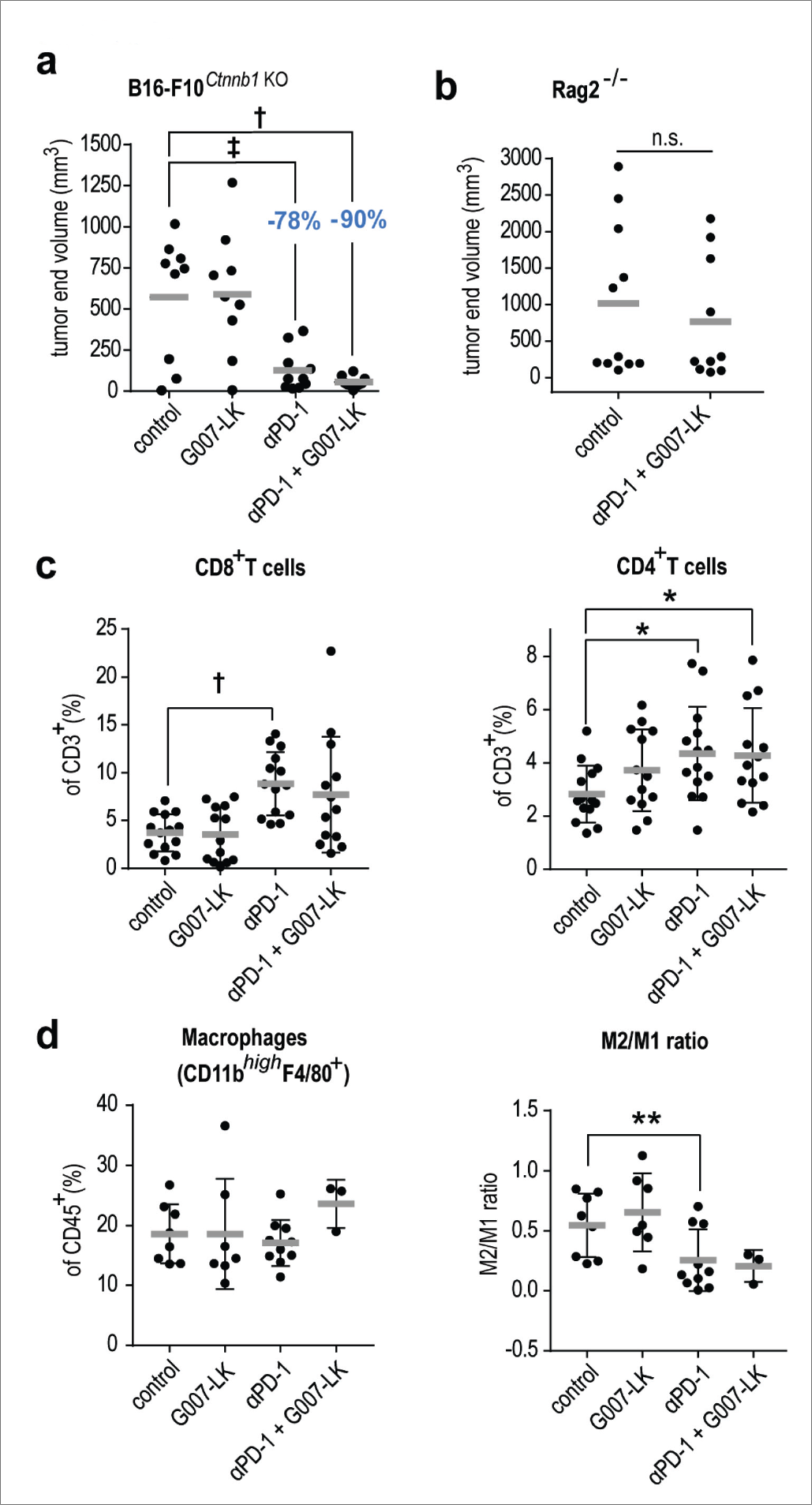
The synergistic tankyrase and anti-PD-1 treatment effect is β-catenin and T cell dependent. **a**, B16-F10^*Ctnnb1*KO^ tumor (s.c.) end volume reduction upon anti-PD-1 (−78%) and combined anti-PD-1/G007-LK treatment (−90%) in mice treated from day 11 through day 25. Control diet (*n* = 9), G007-LK diet (*n* = 9), anti-PD-1 (*n* = 10) and anti-PD-1/G007-LK (*n* = 10). Mann-Whitney rank sum tests are indicated by ^‡^ (*P* < 0.05) and ^†^ (*P* < 0.01). For **a-b**: Mean values are indicated by grey lines. **b**, Tumor end volumes of B16-F10-challenged (s.c.) C57BL/6 Rag2^−/−^ mice treated for 12 days with control diet (*n* = 11) compared to combined anti-PD-1/G007-LK treatment (*n* = 10). For **c** and **d**, flow cytometry analysis of single cell suspensions of comparable-sized tumors collected from B16-F10-challenged (s.c.) C57BL/6N mice treated for 7-17 days with control diet (*n* = 14), G007-LK diet (*n* = 13), anti-PD-1 (*n* = 14) or anti-PD-1/G007-LK (*n* = 13). Mean values are indicated by grey lines ± s.d‥ **c**, T cells (CD3^+^) shown as % of leukocytes (CD45^+^). **c,** CD8^+^T cells and CD4^+^T cells shown as % of CD3^+^ T cells. ANOVA on ranks (Dunn’s method versus control) is indicated by ^†^ (*P* < 0.05) and one-way ANOVA tests (Holm-Sidak method vs. control) are indicated by * (*P* < 0.05). **d**, Macrophages (CD11b^high^F4/80^+^) plotted as % of leukocytes (CD45^+^), and ratio of M2-like (MHCII^*low*^CD206^*high*^) to M1-like (MHCII^*high*^CD206^*low*^) macrophages (M2/M1 ratio). Tumor samples were collected from B16-F10-challenged (s.c.) C57BL/6N mice after 11 days treatment with control diet (*n* = 8), G007-LK diet (*n* = 7), anti-PD-1 (*n* = 10) or anti-PD-1/G007-LK (*n* = 3). Two-tailed t-test is indicated by ** (*P* < 0.05).

Previous work has demonstrated that decreased WNT/β-catenin signaling, in a genetically modified mouse melanoma model, promoted adaptive immune responses within the tumor by enhanced secretion of CCL4^6,19^. The subsequent attraction of dendritic cells to the tumor site supported infiltration and activation of tumor-reactive CD8^+^ T cells^6^. Dependency on T cells for the anti-PD-1/G007-LK synergy was evaluated using B16-F10-challenged CD8^+^ and CD4^+^ T cell-deficient Rag2^−/−^ mice^20^. No significant effect of anti-PD-1/G007-LK treatment was observed in Rag2^−/−^ mice, showing that the presence of T cells is required for the treatment effect (Fig. 2b and Supplementary Fig. 8). To assess immune cell infiltration upon treatment, we performed flow cytometry analysis using tumors of comparable size collected on day 7-17 (Supplementary Fig. 9a and 10). A trend towards increased abundance of leukocytes was observed in the anti-PD-1-treated group, but not between the controls and the G007-LK or anti-PD-1/G007-LK-treated mice (Supplementary Fig. 9b). An increase in total T cell and CD8^+^ T cell infiltration was seen in both the anti-PD-1 and anti-PD-1/G007-LK groups, whereas CD4^+^ T cells were similarly increased in all treatment groups (Fig. 2c and Supplementary Fig. 9c). No discernible differences in the abundance of CD4^+^ or CD8^+^ T cells expressing the memory marker CD44 was observed (Supplementary Fig. 9d). T_regs_ constituted approximately 1% of infiltrating CD45^+^ cells in all groups (Supplementary Fig. 9e). Myeloid DCs were decreased in the anti-PD-1 and anti-PD-1/G007-LK-treated groups while CD103^+^ DCs were equally present in all treatment groups (Supplementary Fig. 9f). Lymphoid DCs, M-MDSC and neutrophils were present at comparable levels across all treatment groups (Supplementary Fig. 9g). A trend towards increased macrophage infiltration was observed in anti-PD-1/G007-LK-treated mice (Fig. 2d). Infiltration of M1-like macrophages was increased and M2-like macrophages decreased in the anti-PD-1 and anti-PD-1/G007-LK-treated groups leading to a decreased M2/M1 ratio (Fig. 2d and Supplementary Fig. 9h). In summary, combined anti-PD-1/G007-LK treatment of B16-F10 tumors confers a general T cell-dependent growth-inhibitory effect on B16-F10 tumors. Changes in myeloid DC, T cell and macrophage infiltrations are likely attributable to anti-PD-1 treatment but do not alone cause anti-tumor activity.

To identify alterations in CCL4 secretion upon G007-LK treatment, conditioned supernatants from matrigel-embedded B16-F10 tumors^21^ and cell culture were screened using ELISA assays. Upon G007-LK treatment, CCL4 secretion was significantly reduced in B16-F10 tumors (Fig. 3a) while no significant CCL4 regulation was seen in cultivated B16-10 cells (Fig. 3b). *Ccl4* expression was not inversely correlated to its previously described negative regulator *Atf3*^*6*^ in neither wild-type B16-F10 cells nor in B16-F10^*Ctnnb1*KO^ cells when compared to wild-type cells (Fig. 3a,b). In conclusion, the anti-PD-1/G007-LK-induced anti-tumor efficacy observed herein is independent from enhanced CCL4 secretion.

**Fig. 3.**
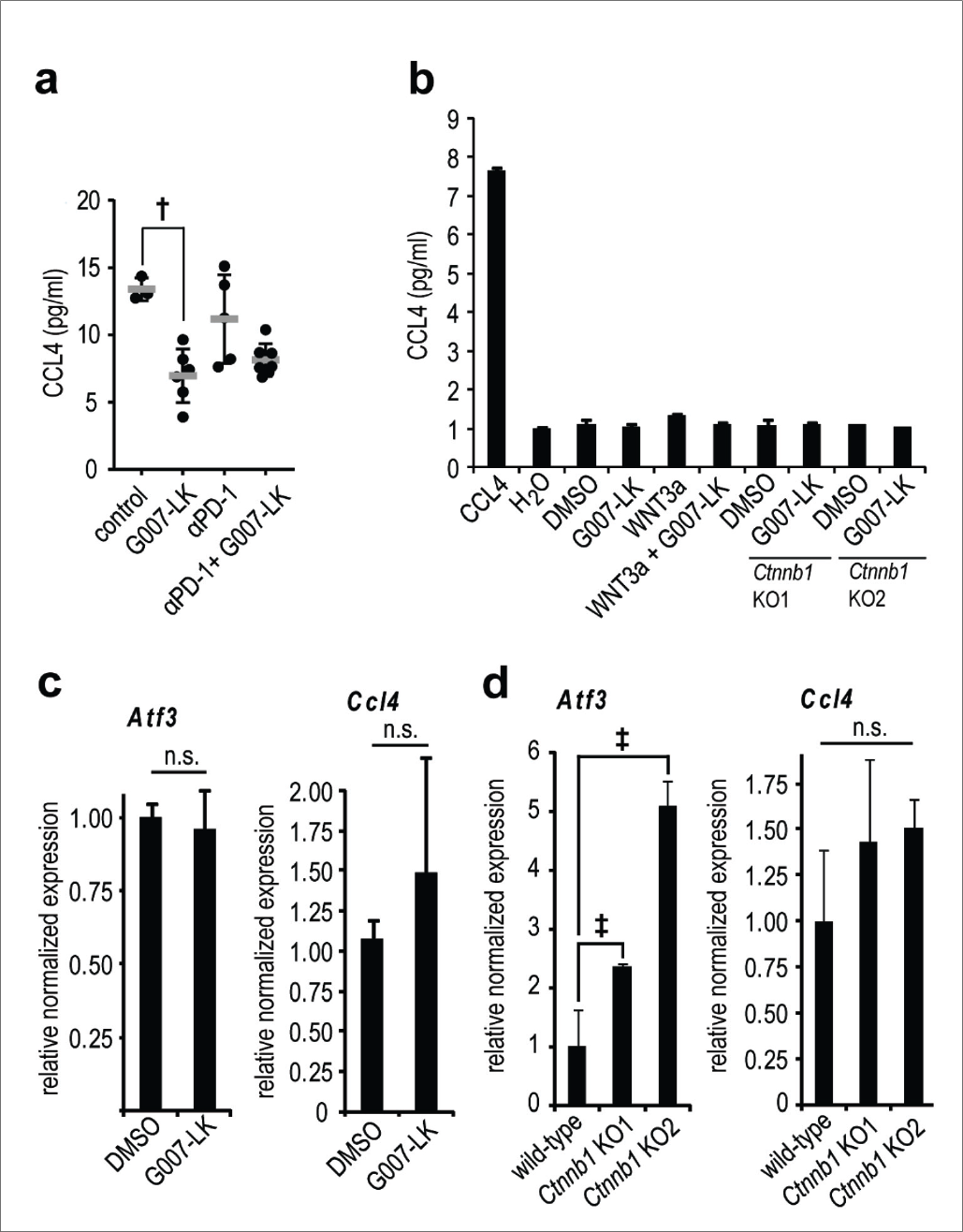
Tankyrase inhibition decreases CCL4 secretion in B16-F10 tumors but not in B16-F10 cell culture. **a**, CCL4 ELISA measurements using conditioned supernatants from matrigel-embedded s.c. B16-F10 tumors screened using multiplex immunoassay upon treatment from day 6 through day 14. Treatments: Control diet (*n* = 3), G007-LK diet (*n* = 6), anti-PD-1 (*n* = 5) and anti-PD-1/G007-LK (*n* = 8). One-way ANOVA on ranks test (Dunn’s method versus control) is indicated by ^†^ (*P* < 0.01). Mean values are indicated by grey lines ± s.d‥ **b**, CCL4 ELISA assay using supernatants from *in vitro* cultured (72 hours) B16-F10 wild-type cells treated with vehicle (DMSO, 0.01%), G007-LK (1 μM), WNT3a, WNT3a + G007-LK as well as B16-F10^*Ctnnb1*KO^ cells (KO1 and 2) treated with G007-LK (1 μM). Positive control: CCL4. Negative control: H_2_O. Mean values ± s.d. from 2 independent experiments are shown. All samples display measured signals similar to negative control. **c**, Real-time RT-qPCR analyses of *Atf3* and *Ccl4* from B16-F10 cell culture treated (24 hours) with vehicle (DMSO, 0.01%) or G007-LK (1 μM). n.s. = not significant. **d**, Real-time RT-qPCR analyses of *Atf3* and *Ccl4* from cultured B16-F10^*Ctnnb1*KO^ cells compared to wild-type B16-F10 cells. Two-tailed Mann-Whitney rank sum tests are indicated by ^‡^ (*P* < 0.05) and n.s. = not significant.

B16-F10 murine melanoma, lacking the BRAF^V600E^ mutation^22,23^, only partially recapitulates the genetic features of human melanoma. Thus, murine B16-F10 cells, and a panel of 18 human melanoma cell lines, were exposed to G007-LK treatment followed by RNA sequencing and bioinformatic analyses. In the untreated group, no clear correlation was found between baseline transcription of WNT/β-catenin signaling target genes or the mutation load, including BRAF^V600E^, against overall gene expression (Supplementary Fig. 11a and 12). However, when the cell lines were classified into two groups based on baseline YAP signaling target gene transcription, YAP^high^ and YAP^low^ (Fig. 4a), the YAP^high^ group coincided with clustering of overall gene expression and against a panel of markers for β-catenin-controlled melanoma cell fate and proliferation^24^ (Supplementary Fig. 11). The markers in the the YAP^high^ group included low relative transcription of *MITF* (*MITF*^low^)(Fig. 4b). MITF is a lineage-restricted regulator in melanocytes that is positively associated with melanoma proliferation, and also suppressed invasion and metastasis^25–27^. Indeed, when using untreated samples, Ingenuity Pathway Analysis (IPA) core analysis of the YAP^high^ versus the YAP^low^ group revealed *MITF* as the most significant key upstream transcriptional regulator separating the two groups (Supplementary Fig. 13 and 14). Upon treatment with G007-LK, samples in the *MITF*^low^ group were predisposed for decreased *MITF* expression (*MITF*^decreased^), while the *MITF*^high^ group was oppositely increased (Fig. 4b and Supplementary Fig. 15a). In B16-F10 tumors, transcription of *Mitf* was reduced upon G007-LK-treatment (Fig. 4c). Attenuated WNT/β-catenin and/or YAP signaling activity was observed in nearly all samples upon G007-LK treatment, but did not correlate with changes in *MITF* expression, nor act as a predictive marker for *MITF* regulation (Fig. 4d,e and Supplementary Fig. 15 and 16). In conclusion, YAP^high^ is a marker for a melanoma sub-group that includes B16-F10 and tracks with tankyrase inhibitor-induced reduction in *MITF* expression (Fig. 4e and Supplementary Fig. 17).

**Fig. 4.**
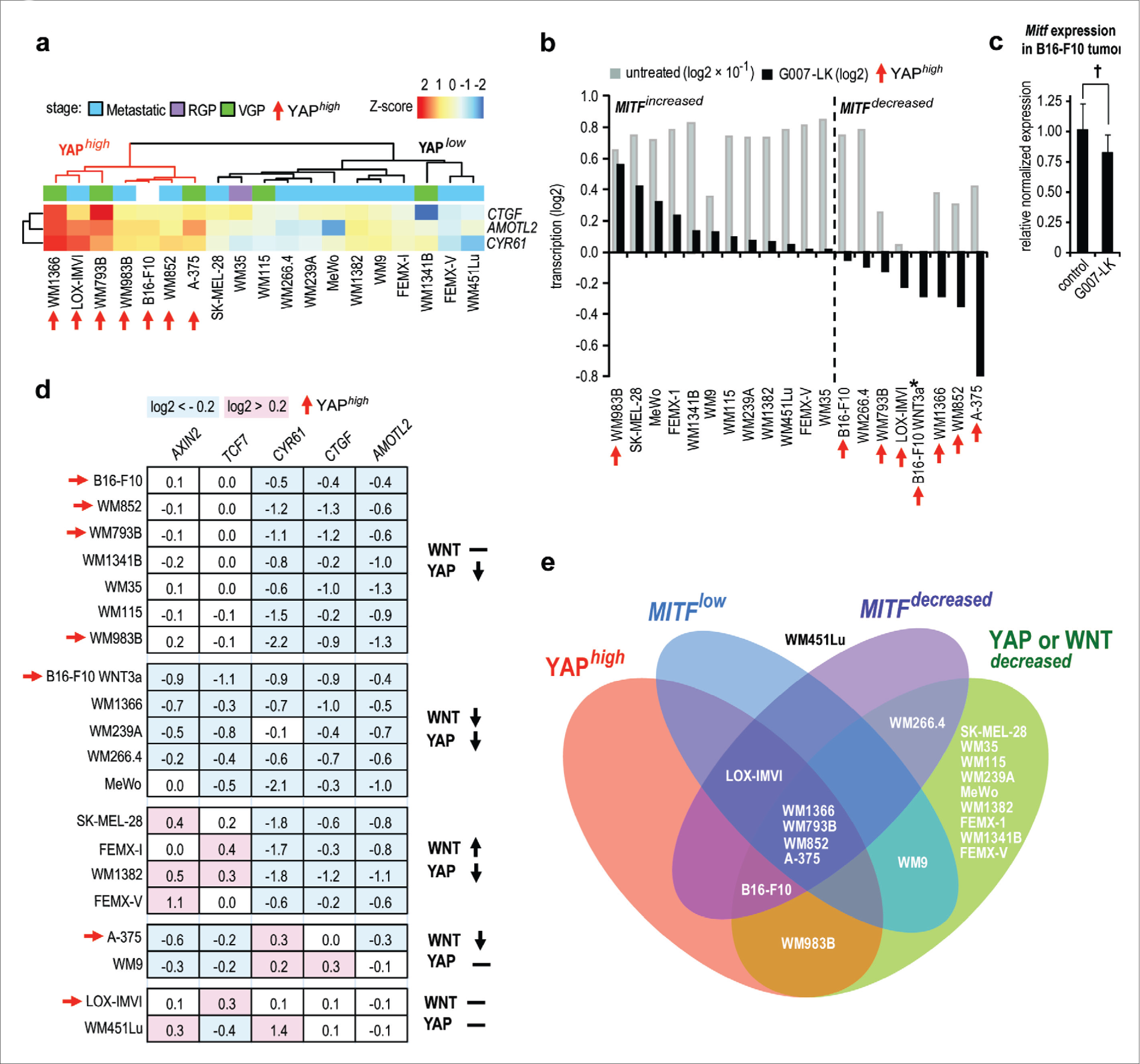
High activity of YAP signaling correlates with low baseline *MITF* expression and potential for decreased *MITF* transcription upon tankyrase inhibition. **a**, Heatmap and clustering of YAP signaling target transcripts (*Cyr61*, *Ctgf* and *Amotl2*) for 18 untreated human melanoma cell lines (pooled triplicates used for RNA sequencing) and murine B16-F10 melanoma (biological triplicates used for RNA sequencing). 7 of 19 samples displayed high relative transcription of YAP signaling target genes (YAP^high^) when compared to samples with less transcription (YAP^low^). YAP^high^ is highlighted by orange branches in the dendrogram and arrows. Scale bar indicates differences in Z-score values for log2 transcripts per millions (TPMs) within each row. Cancer stages: Metastatic in blue, radial growth phase (RGP) in pink and vertical growth phase (VGP) in green. **b**, Bar chart showing *MITF* expression in untreated samples (grey bars, log2-transformed TPMs × 10^−1^) and upon treatment with G007-LK (1 μM) for 24 hours (black bars sorted descending from left to right, log2 values from treated versus untreated TPMs). The dotted vertical line depicts division between samples with increased (*MITF*^increased^, left) or decreased (*MITF*^decreased^, right) expression of *MITF* upon tankyrase inhibitor treatment. * indicates that no value for untreated sample is inserted. For **b** and **d**: B16-F10 WNT3a = WNT3a + G007-LK relative to WNT3a-stimulated control. YAP^high^ cell lines are highlighted by orange arrows. **c**, Real-time RT-qPCR analysis of *Mitf* expression in G007-LK-treated B16-F10 s.c. tumors versus control tumors (*n* = 8). Two-tailed Mann-Whitney rank sum test is indicated by ^†^ (*P* < 0.01). Mean values ± s.d. for 2 repeated measurements are shown. **d**, Changes in gene expression for WNT/β-catenin (*Axin2* and *Tcf7*) and YAP (*Cyr61*, *Ctgf* and *Amotl2*) signaling target genes for 18 G007-LK-treated (1 μM) human and murine B16-F10 melanoma cell lines. Down-regulated signaling activity is indicated by ↓ (log2 < −0.2, highlighted in blue), up-regulated activity is indicated by ↑ (log2 > 0.2, highlighted in pink), and inconclusive or lack of regulation is indicated by **−**. **e**, Venn diagram depicting the cell lines intersecting profiles for pre-treatment (YAP^high^ and *MITF*^low^) and response to G007-LK treatment (*MITF*^decreased^ and YAP or WNT^decreased^).

WNT/β-catenin signaling plays a central role, not only in regulation of immune cell homeostasis, development and function, but also impacts circulating immune cells, - including control of peripheral T cell activation, differentiation and tumor-immune cell interplay^19^. Active YAP signaling may support immune evasion in cancer and melanoma by inducing PD-L1 expression^28,29^ (Supplementary Fig. 18) while another report shows that enhanced YAP signaling, due to loss of LATS1/2, may promote anti-cancer immune response^30^. Tankyrase inhibition also leads to regulation of the complex network of cytokines and chemokines, which conceivably may influence myeloid cell anti-tumor activity^31^ (Fig. 3 and Supplementary Fig. 19). Presently, neither the meeting points between MITF, its regulation by YAP/TAZ/TEAD, AP-1 and tankyrase, nor the function in immune regulation and control of susceptibility to checkpoint-inhibitor therapy are well characterized^27,32–36^. Collectively, the results presented herein suggest that G007-LK-induced blockade of WNT/β-catenin and YAP signaling pathways affects a complex network of cytokines and chemokines leading to improved efficacy of PD-1 immune checkpoint blockade. The findings rationalize a further in-depth preclinical and clinical evaluation of combining checkpoint inhibitors with tankyrase inhibition for treatment of melanoma.

## Methods

### Cell culture

The mouse melanoma cell line B16-F10 was obtained from the American Type Culture Collection (ATCC® CRL-6475™). Clone M-3^Z1^ (ProQinase) cells were derived from Clone M-3 cells (ATCC^®^ CCL-53.1^™^) after implantation in DBA/2N mice. The cell cultures were kept at low-passage numbers in RPMI-1640 medium (R8758, Sigma Aldrich). The cell culture medium was supplemented with 1% Penicillin-Streptomycin (P4333, Sigma Aldrich) and 5% Fetal Bovine Serum (FBS, 10270-106, Gibco) and grown at 37°C in humidified incubators with 5% CO_2_. The cells were routinely monitored for mycoplasma using MycoAlert Mycoplasma detection kit (Lonza). B16-F10 cells were authenticated by short tandem repeat profiling and subsequent analysis confirming C57BL/6 origin (Leibniz-Institute DSMZ). General protocol for treatment of cultured B16-F10 cells: Cells were seeded one day before treatment to reach ~80% confluence for 24 hours treatments and ~20% confluence for 72 hour treatments. The cell culture medium was changed for medium containing 0.01% dimethyl sulfoxide (DMSO, D8418, Sigma Aldrich), 1 μM or various doses of G007-LK (Mercachem or ChemRoyal), 0.5 μg/mL WNT3a (Recombinant Mouse Wnt3a, 1324-WN-010, R&D Systems) or a combination of G007-LK (1 μM) and WNT3a (0.5 μg/mL).

### CRISPR/Cas9-based knock-out

Genomic RNA sequences targeting exon 4 of mouse *Ctnnb1* were designed using the web-based CHOPCHOP platform^37^: *Ctnnb1*_gRNA1: 5’-GATTAACTATCAGGATGACG-3’. The gRNA sequence was inserted in to the pSPCas9(BB)-2A-GFP vector (PX458, 48138, Addgene, provided by Dr. Feng Zhang^38^), and the vector was delivered to tumor cells by transfection using polyethylenimine (408727, Sigma Aldrich). After 24 hours, GFP-expressing tumor cells were single-cell sorted (BD FACSAria II, BD Biosciences). Genomic DNA was isolated from individual clones, the relevant gene fragment was amplified by PCR (Forward primer: 5’-GTTCCCTGAGACGCTAGATGAG-3’. Reverse primer: 5‘-ACATCACTGCTTACCTGGTCCT-3’.) and screened by Sanger sequencing for non-sense mutations in *Ctnnb1*. Additional verification of gene knockout was performed by immunofluorescence staining (primary antibody β-catenin [BD610153, 1:500, BD Biosciences] and secondary antibody goat anti-mouse IgG Alexa 488 [A28175, 1:500, ThermoFisher]) and Western blot analysis (β-catenin, 1:500).

### Western blot analysis

Treated cells were washed in PBS and lysed in NP40 Cell lysis buffer (Invitrogen) containing protease inhibitors. The nuclei were pelleted and separated from the cytoplasmic supernatant fractions. RIPA lysis buffer containing phosphatase and protease inhibitors was added to the nuclei followed by sonication (Bioruptor^®^Plus, Diagenode). Protein concentrations were measured using Pierce™ BCA Protein Assay Kit (Pierce Biotechnology). The protein samples were separated by SDS PAGE gel electrophoresis (Invitrogen) and immunoblotted (Immobilon-PSQ PVDF Membrane, Millipore) using the following primary antibodies: Tankyrase-1/2 (TNKS1/2, H-350, sc-8337, Santa Cruz Biotechnology), non-phospho (active) β-catenin (D13A1, 8814, Cell Signaling Technology), total β-catenin (610153, BD Transduction Laboratories™), TAZ (HPA007415, Sigma Aldrich), YAP (sc-101199, Santa Cruz Biotechnology), phospho-YAP (Ser127, 4911, Cell Signaling Technology), GSK3β (12456, Cell Signaling Technology), phospho-GSK3β (Ser9) (9323, Cell Signaling Technology). GAPDH (sc-32233, Santa Cruz Biotechnology), β-Tubulin III (T2200, Sigma Aldrich), actin (A2066, Sigma Aldrich) and lamin B1 (ab16048, Abcam) were used as loading controls. Primary antibodies were visualized with secondary HRP-conjugated antibodies (mouse anti-rabbit IgG-HRP, sc-2357, Santa Cruz Technology or donkey anti-rabbit IgG, 711-035-152, Jackson ImmunoResearch Inc.) and enhanced with chemiluminescent substrate (ECL™ Prime Western Blotting Detection Reagent, RPN2236, GE Healthcare) and ChemiDoc™ Touch Imaging System (Bio-Rad).

### RNA isolation and real-time qRT-PCR

Total RNA was isolated from treated cell lines and tumor samples using GenEluteTM Mammalian Total RNA Miniprep Kit (Sigma Aldrich). The RNA concentration was measured using Nanodrop 2000c spectrophotometer (Thermo Scientific). cDNA was synthesized from the purified RNA using SuperScript™ VILO cDNA Synthesis Kit (Invitrogen). Real-time qRT-PCR (TaqMan®Gene Expression system, Applied Biosystems) was performed using Viia7 (Applied Biosystems). The following probes were used (all from Applied Biosystems): *Axin2* (Mm00443610_m1), *Tcf7* (Mm00493445_m1), *Atf3* (Mm00476033_m1), *Ccl4* (Mm00443111_m1), *Amotl2* (Mm00502287_m1), *Ctgf* (Mm01192933_g1) and *Cyr61* (Mm00487498_m1).

### Proliferation assays

1000 cells/well were seeded in 96-well plates using minimum 6 replicates for each treatment tested. The day after, cell culture medium was changed to contain various doses of G007-LK or vehicle (DMSO, Sigma Aldrich) and the plates were incubated in an IncuCyte (FLR30140, Essen BioScience) at 37 °C for real-time monitoring of cell confluency. At experiment endpoint (80-100% confluency after 5-7 days of cell growth), the cells were incubated for 1 hour at 37 °C with CellTiter 96® AQueous Non-Radioactive Cell Proliferation Assay (MTS, Promega) according to the supplier’s recommendations. Abs_490_ was measured spectrophotometrically (Wallac 1420 Victor2 Microplate Reader, Perkin Elmer) and compared to Abs_490_ (t0) using the following formula to determine single well values relative to the DMSO vehicle control: (sample A_490_ − average A_490 t0_)×100/ (average A_490_ [for 0.01% DMSO controls] − average A_490 t0_).

### Immunofluorescent staining and Structured Illumination Microscopy (SIM)

Cells grown on coverslips pre-coated with poly-L-lysine (sc-286689, Santa Cruz Biotechnology) were fixed in 4% paraformaldehyde (P6148, Sigma Aldrich, 15 minutes at room temperature) and permeabilized with 0.1% Triton-X100/PBS (T8787, Sigma Aldrich, 15 minutes at room temperature) followed by 1 hour incubations with primary and secondary antibodies diluted in PBS with 4% bovine serum albumin, both at room temperature. Nuclear counterstaining was performed with DAPI (D9542, Sigma Aldrich, 1 μg/mL, 5 minutes at room temperature) and coverslips were mounted in ProLong Diamond Antifade Mountant (Thermo Fisher Scientific). The following primary antibodies and dilutions were used: β-catenin (BD610153, 1:500, BD Biosciences) and Tankyrase-1/2 (E10, sc-365897, 1:50, Santa Cruz Biotechnology). Secondary antibodies used (both from Thermo Fisher Scientific, 1:500): Anti-rabbit IgG Alexa488 (A-21206) and anti-Mouse IgG Alexa594 (A-11005). SIM images were acquired on a Zeiss Elyra PS1 microscope system using standard filters sets and laser lines with a Plan-APOCHROMAT 63x 1.4 NA oil objective. SIM imaging was performed using 5 grid rotations with the 0.51 μm grid for 20 Z planes with a Z-spacing of 0.184 nm between planes. SIM images were reconstructed with the following “Method” parameters in the ZEN black software (MicroImaging, Carl Zeiss): Processing: Manual, Noise Filter: −5, SR Frequency Weighting: 1, Baseline Cut, Sectioning: 100/83/83, Output: SR-SIM, PSF: Theoretical. The SIM images are displayed as maximum intensity projections rendered from all Z planes.

### Luciferase reporter assays

The following plasmids were used: SuperTOP-luciferase (WNT/β-catenin signaling pathway reporter with 7X TCF binding sites: ST-Luc, gift from V. Korinek), SFF-Luc (negative control reporter with mutated TCF binding sites: SuperFOPflash-luciferase, gift from V. Korinek), 8xGTIIC-luciferase (hippo and YAP signaling pathway reporter: 34615, Addgene, provided by Dr. Stefano Piccolo^39^) and *Renilla* luciferase (pRL-TK, Promega). On day 1, B16-F10 cells were seeded in 10-cm dishes to reach 50-60 % confluency for co-transfections on day 2 (16.5 μg luciferase reporter and 3 μg Renilla luciferase, FuGENE*®* HD, Promega). On day 3, the cells were trypsinized and seeded in 96-well plates and treatment was added on day 4. On day 5, the cells were lysed and the firefly and *Renilla* luciferase activities were measured using Dual-Luciferase Reporter Assay (Promega) protocol and GloMax®-Multi Detection System (Promega). XLfit (Idbs) was used to calculate the half maximal inhibitory concentration (IC_50_) using the Langmuir Binding Isotherm formula: fit = ((A+(B×x)) + (((C-B)×(1-exp ((−1×D)×x)))/D)), res = (y-fit).

### Tumor initiation, treatment and analysis of mouse melanoma *in vivo*

General procedure: Subcutaneous B16-F10 (B16-F10, ATCC® CRL-6475™) and B16-F10^*Ctnnb1*KO^ tumors (using KO clone 1) were implanted (left flank, 0.2×10^6^ tumor cells in 20 μl PBS) in female B6N-Tyr^c-Brd^/BrdCrlCrl albino mice (C57BL/6N, Charles River) or female RAGN12F; B6.129S6-Rag2^tm1Fwa^ N12 mice (Rag2 [C57BL/6 background], Taconic Biosciences). 1.0×10^6^ Clone M-3(Z1) cells (ProQinase) in 100 μl were subcutaneously implanted in female DBA/2NCrl (DBA/2N mice (Charles River). All mice were between 3-8 weeks old at experiment startup. Randomization of mice to create treatment groups with ~equal tumor sizes was performed by random number generation within individual blocks (MS-Excel 2016). Treatments used: i) Control (Purina 5001 diet, Research Diets Inc), ii) G007-LK (250 mg/kg in diet [ab libitium] that delivers 43-53 mg/kg/day, depending on measured [3×weekly] consumed diet [3.2-4.4 g/mouse/day] and body weight), iii) anti-PD-1 (10 mg/kg, intraperitoneally, i.p. [RMP1-14, BE0146, batch 614616A2, Bio X Cell]), iv) anti-PD-L1 (10 mg/kg, i.p. [10F.9G2, batch 6154598816S1, Bio X Cell]), v) combinations of G007-LK and anti-PD-1 or vi) G007-LK and anti-PD-L1. Previous analyses have documented the pharmacokinetics of G007-LK in female mice using oral gavage delivery (t_1/2_ = 2.6 hours and bioavailability = 76%) and in diet containing 250 mg/kg G007-LK (t_1/2_ = 4.2 hours)^7,40^. Compound delivery was monitored via food consumption. The chow was weighed thrice weekly to calculate G007-LK delivery per animal and day. For experiments using B16-F10 *Ctnnb1* knock-out cells in C57BL/6N mice: Intraperitoneal injections of anti-PD-1 were administered on day 11, 15 and 18. For experiments using B16-F10 cells in C57BL/6N mice: Intraperitoneal injections of anti-PD-1 or anti-PD-L1 were administered on day 10, 13 and 17, -and every 3-4 days from day 21 until the end for the survival analysis. For experiments in Rag2 mice or for Clone M3(Z1) in DBA/2N mice: Intraperitoneal injections of anti-PD-1 were administered on day 8, 11 and 15. Primary tumors were measured by caliper (OMC Fontana) and tumor sizes calculated according to the formula W^2^×L/2 (L= length and W= width).

Tumor growth assay using B16-F10 *Ctnnb1* knock-out cells in C57BL/6N mice: On day 11, 48 tumor-bearing animals (primary tumors reaching 20-100 mm^3^) were randomized into 4 groups (n = 12), each treated with i), ii), iii) or v). On day 25, mice were euthanized, and the tumors were dissected, weighed and volumes re-measured with caliper. 3 mice from treatment group i) and 2 mice from treatment groups ii), iii) and v) were euthanized for ethical reasons before experiment termination, due to skin ulcerations in the tumor area or excessive tumor size.

Assay for the efficacy of G007-LK-mediated inhibition of tankyrase and WNT/β-catenin and YAP signaling pathways using B16-F10 cells in C57BL/6N mice: Mice with primary tumor volumes of 20-40 mm^3^ were randomized into two groups (control or G007-LK for 4 nights, n = 8). Whole tumor protein extracts were prepared by sonication in an ultrasound bath (Bioruptor^®^ Plus, Diagenode) in RIPA buffer (Thermo Fisher Scientific) containing phosphatase and protease inhibitors (PhosStop, 4906837001 and cOmplete™ Protease Inhibitor Cocktail, 4693116001, both from Roche). Tumor tissues were homogenized using MagNA Lyser Green Beads (Roche) and total mRNA isolated using GenEluteTM Mammalian Total RNA Miniprep Kit (Sigma Aldrich).

Tumor growth assay using B16-F10 cells in C57BL/6N mice: On day 10, 72 tumor-bearing animals (primary tumors reaching 20-100 mm^3^) were randomized into 6 groups (n = 12), and treatments i) - vi) were administered to the mice. On day 21, the experiment was terminated, and tumors from euthanized mice were dissected, weighed and volumes re-measured with caliper. For 6 euthanized animals from treatment groups i), ii), iii) and v): small intestinal was dissected, formalin-fixed (10%), paraffin-embedded in a Swiss roll configuration and sections were stained with H&E (115938 and 115935, Merck Millipore). 1, 2, 1, 4, 3 and 2 mice from treatment groups i), ii), iii), iv) and v), respectively, were found dead or euthanized for ethical reasons during the experiment due to anemia or skin ulcerations in the tumor area.

Clone M-3(Z1) tumor growth assay: On day 8, 48 tumor-bearing animals (with primary tumors reaching approximately 26 mm^3^) were randomized into 4 groups (n = 12) and treatments i), ii), iii) and v) were administered to the respective mice. On day 18, the animals were euthanized and the tumors were dissected, weighed and volumes re-measured with caliper. 4, 1, 3 and 1 mice from treatment groups i), ii), iii) and v), respectively, were euthanized for ethical reasons during the experiment due to skin ulcerations in the tumor area.

Tumor growth assay using B16-F10 cells in B6.129S6-Rag2tm1Fwa N12 mice (Rag2^−/−^)^20^ mice: On day 8, 24 tumor-bearing animals (with primary tumors reaching 18-61mm^3^) were randomized into 2 groups (n = 12), and treatments i) and v) were administered to the respective mice. On day 20, the animals were euthanized. 1 mouse from treatment group i) and 2 mice from treatment group v) were euthanized for ethical reasons before experiment termination, due to skin ulcerations in the tumor area or excessive tumor size.

Survival assay using B16-F10 cells in C57BL/6N mice: 60 animals with primary tumors reaching 20-100 mm^3^ were randomized into 2 groups (n = 30, control or G007-LK and anti-PD-1). The survival assay endpoint criterion was set to tumor volume >1000 mm^3^. 11 mice in the control group and 14 mice in the treatment group were euthanized (for ethical reasons) or found dead due to anemia or skin ulcerations in the tumor area during the experiment before reaching the endpoint criterion. Tumors were collected from surviving and treated animals at endpoint on day 38, and from control animals euthanized on day 26. The tumors were dissected, fixed (10% formalin), embedded in paraffin, sectioned and stained with H&E (115938 and 115935, Merck Millipore). For immunostaining 2.5 μm sections were boiled for 20 minutes in 10 mM citrate buffer pH 6.0 and incubated with primary antibody F4/80 (2 μg/mL, clone CI:A3, ab6640, Abcam) in PBS with 1.25% BSA overnight at 4°C, and then incubated with fluorescently labeled secondary antibody (donkey anti rat IgG Alexa Fluor 488, 5 μg/mL, Molecular Probes) for 60 minutes at 37°C. Hoechst 33258 nuclear dye (0.5 μg/mL, Sigma Aldrich) was added to the final washing solution. Pictures were captured using a Nikon Eclipse model N *i*-U microscope (Nikon) equipped with Nikon Plan-Fluor objective lenses and an Infinity 2 digital camera (Lumenera Corporation).

All animal experiment described in this paragraph were performed by ProQinase GmbH, following approval by local animal experiment authorities (Freiburg, Germany) and in compliance with FELASA guidelines and recommendations.

### Tumor flow cytometry analysis

Primary tumor from treatment groups i), ii), iii) and v) was collected and processed for flow cytometry analysis to determine the presence of sub-populations of T cells, myeloid-derived suppressor cells and macrophages (carried out by ProQinase). For analysis of T cells and myeloid-derived suppressor cells, animals were treated for 7-17 days to obtain similarly distributed primary tumor volumes ranging from 80-240 mm^3^ (Supplementary Fig. 9a). For analysis of macrophages, tumor material obtained at endpoint of the tumor growth assay in C57BL/6N mice was collected on day 21. Both animal experiment were performed by ProQinase GmbH, following approval by local animal experiment authorities (Freiburg, Germany) and in compliance with FELASA guidelines and recommendations. Tumors were disrupted using gentleMACS Tubes (Miltenyi Biotec) containing the enzyme mix of the Tumor Dissociation Kit according to the manufacturer instructions (Miltenyi Biotech). Erythrocytes were removed with the Red Blood Cell Lysis Solution (Miltenyi Biotech). Single cell suspensions were counted, and up to 3×10^6^ cells/well were dispensed into 96-well plates. The single cells were washed with PBS and stained for living cells (eBioscience™ Fixable Viability Dye eFluor™ 455UV [65-0868-14, eBioscience]). After washing and centrifugation (400 x *g*), the samples were incubated with 50 μl/well of Fc block (anti-mouse CD16/CD32, 1:50, 14-0161-85, clone 93, eBioscience) for 30 minutes in FACS buffer (PBS with 2% FCS and 0.2% EDTA, 03690, Sigma Aldrich). Thereafter, 50 μl of the following 2X concentrated antibody master mixes were added to each well and incubated for 30 minutes in the dark. For T cells, the following antibodies against murine targets were used: CD45 (CD45-PacBlue [30-F11], 48-0451-82, eBioscience), CD3 (CD3-Violet 605 [17A2], 100237, BioLegend), CD4 (CD4-APC-Cy7 [GK1.5], 47-0041-82, eBioscience), CD8a (CD8-PerCP [53-6.7], 553036, BD Pharmigen), CD25 (CD25-APC [PC61.5], 17-0251-82, eBioscience) and CD44 (CD44-FITC [IM7], 11-0441-82, eBioscience). For myeloid cells, the following antibodies against murine targets were used: CD45 (CD45-PacBlue [30-F11], 48-0451-82, eBioscience), CD11b (CD11b-FITC [M1/70], 11-0112-82, eBioscience), CD11c (CD11c-APC-Cy7 [N418], 47-0114-80, eBioscience), Ly6G (Ly6G-APC [RB6-8C5], 17-5931-81, eBioscience), Ly6C (Ly6C-PE [HK1.4], 12-5932-82, eBioscience) and CD103 (CD103 [Integrin alpha E], PerCP-Cy5.5 [2E7], 121415, BioLegend). For macrophages, the following antibodies against murine targets were used: CD45 (CD45-PacBlue [30-F11], 48-0451-82, eBioscience), CD11b (CD11b-FITC [M1/70], 11-0112-82, eBioscience), F4/80 (F4/80-APC-Cy7 [BM8], 47-4801-82, eBioscience), CD206 (CD206-PE [C068C2], 141706, BioLegend) and MHC II (MHC II-APC [M5/114.15.2], 17-5321-82, eBioscience). After washing, cells were stained for myeloid and macrophage markers and fixed for flow cytometry analysis. Cells stained with the T cell panel markers were prepared for intracellular staining by adding 50 μl fixation/permeabilization (1:3) buffer (00-5523-00, eBioscience) for 30 minutes. Thereafter, 100 μl 1X permeabilization buffer (00-5523-00, eBioscience) was added and the cells were centrifuged at 400 x *g*. The cell pellet was resuspended in 1X permeabilization buffer containing anti-FoxP3 antibody (FoxP3-PE [FJK-16s], 12-5773-82, eBioscience) and incubated for 30 minutes in the dark. After washing twice with permeabilization buffer, the cells were washed with FACS buffer. Cells were kept at 4°C in the dark until analysis. The samples were analyzed by flow cytometry using an LSR Fortessa (Beckton Dickenson) and the gating is shown in Supplementary Fig. 10.

### ELISA

Cell supernatants from treated cells and conditioned B16-F10 tumor supernatants (see description for multiplex immunoassay) were assayed using Mouse CCL4/MIP-1 β Quantikine ELISA Kit (MMB00, R&D Systems) according to the manufacturers’ protocol.

### RNA sequencing

The following samples were used for RNA sequencing experiments and RNA was isolated using GenEluteTM Mammalian Total RNA Miniprep Kit (Sigma Aldrich): Three biological replicates of cultured B16-F10 cells, treated with DMSO (0.01%), G007-LK (1 μM), WNT3a (0.5 μg/mL) or G007-LK and WNT3a for 24 hours. And in addition, three pooled technical replicates from a panel of 18 human melanoma cell lines treated with DMSO (0.01%) or G007-LK (1 μM) for 24 hours: The cell lines SK-MEL-28, MeWo, and A-375 were obtained from the American Type Culture Collection (ATCC). WM35, WM115, WM1341B, WM1366, WM983B, WM451Lu, WM239A, WM266.4, WM852, WM1382, WM9, WM793B were obtained from the Wistar Institute. LOX-IMVI, FEMX-I and FEMX-V were established at the Norwegian Radium Hospital (Oslo, Norway). Cell line authentication was performed by short tandem repeat profiling and subsequent analysis at the Norwegian Radium Hospital. RNA sequencing (TruSeq Stranded mRNA kit for library prep and NextSeq500 v2 chemistry used for sequencing, both Illumina Inc.) was performed at the Genomics Core Facility Oslo (Oslo University Hospital, Norway).

### DNA sequencing

Kinome targeted re-sequencing of the 18 human melanoma cell lines was performed using the SureSelect Human Kinome kit (Agilent Technologies), with capture probes targeting 3.2 Mb of the human genome, including exons and untranslated regions (UTRs) of all known kinases and selected cancer-related genes (to a total of 612 genes). Library construction and in solution capturing was performed following Agilent’s SureSelectXT library construction kit and SureSelect Target enrichment protocol, respectively. Sequencing was performed on an Illumina HiSeq2500 using the TruSeq SBS Kit v5 generating paired-end reads of 75 bp in length. Base calling, de-multiplexing and quality filtering was performed using Illumina’s software packages SCS2.8/RTA1.8 and Off-line Basecaller-v1.8.

### Bioinformatics

Transcripts were quantified with kallisto (v0.44)^41^ using ensembl transcriptome release 91 for human (GRCh38) and release 92 for mouse (GRCm38)^42^. Ensemble BioMart was used to map human orthologs in mouse^41^. Differentially expressed genes (DEGs) were identified with sleuth (v0.29)^43^, limma^44^ (did not result in any comparisons with adjusted P values < 0.05) and DESeq2^45^ in the R programming environment (The R Project for Statistical Computing). The R-package NMF^46^ was used to make hierarchical clusters with TPM values as input. For detection of probable driver mutations, the RNASeq data was aligned with HISAT2 (v2.1.0)^47^ before VarDict (v1.2)^48^ restricted to SNVs reported more than once in COSMIC (v82)^49^ was applied. Due to the variable coverage in RNASeq data, additional SNVs found in an external unpublished gene panel sequencing experiment for the same cell lines, were included. Variants found in both data sets were annotated using ANNOVAR^50^. The scripts used to process and analyze the data and make the related figures and tables are available at https://github.com/ous-uio-bioinfo-core/waaler-et-al-2018 (available to reviewers upon request). The data analysis was performed by the Bioinformatics Core Facility (Oslo University Hospital, Norway). DEGs analysis data, including log2-fold change and adjusted p-values, were uploaded into Ingenuity Pathway Analysis (IPA) version 01-10 (Qiagen). The DEGs analysis data was analyzed using the core analysis function with the Ingenuity Knowledge Base (genes only) reference set and direct relationships, with no filters set for node types, data sources, confidence, species, tissues & cell lines and mutations. For the IPA core analyses, the log2fold and or adjusted p-value cutoffs are specified in the figure legends.

### Statistical analyses

Sigma Plot® 12.5 (Systat Software Inc.) was used to perform statistical tests: ANOVA or Student’s t-test for comparisons with homogeneous variances (Shapiro-Wilk test, P > 0.05) and ANOVA on the ranks or Mann-Whitney rank sum tests for comparisons where the normality assumption was violated (Shapiro-Wilk test, P < 0.05). GrapPad Prism 7 was used for Kaplan-Meyer estimations and statistical analysis. Single outlier detections were identified by Dixon’s and/or Grubb’s tests (threshold, P < 0.05) using ControlFreak (Contchart software).

### Data availability

The RNA sequencing data for this study is available from ArrayExpress (available to reviewers upon request) with accession xxx for the mouse experiment (fastq and processed) and with accession xxx for the human experiment (processed). Supplementary Figures 1-19 are available online. Any additional data generated and analyzed in this study are available from the corresponding author upon reasonable request.

## Acknowledgements

J.W., L.M., NT.S., PA.O., and S.K. were supported by the Research Council of Norway (grant no. 262613, 267639), by South-Eastern Norway Regional Health Authority (grant no. 16/00528-9, 15/00779-2 and 2015012) and from the Norwegian Cancer Society (grant no. 5803958). A.T. was supported by grants from the Norwegian Cancer Society (grant no. 189562 and 181674). A.C. was supported by funding from the Research Council of Norway (grant no. 262814).

## Author Contributions

J.W., L.M., A.T. and S.K. conceived the project, designed the general study, designed animal experiments and interpreted results. J.W., L.M., A.T, NT.S., P.A.O., K.L., S.A.B., M.L. and E.D. performed and analyzed *in vitro* experiments. J.W., L.M. and A.T. performed multiplex immunoassay. S.L.B and KM.H. prepared samples for RNA sequencing. M.F.S. and V.N. performed bioinformatics analyses. E.H. provided DNA sequencing data and support for bioinformatics analysis. C.H. performed IHC and analysis. A.C. provided support for IHC and analysis. J.W., L.M., A.T. and S.K. wrote the manuscript with feedback from all authors.

