## Supplementary Figures and Legends for "Tankyrase inhibition sensitizes melanoma to PD-1 immune checkpoint blockade in syngeneic mouse models"

#### Supplementary Fig. 1

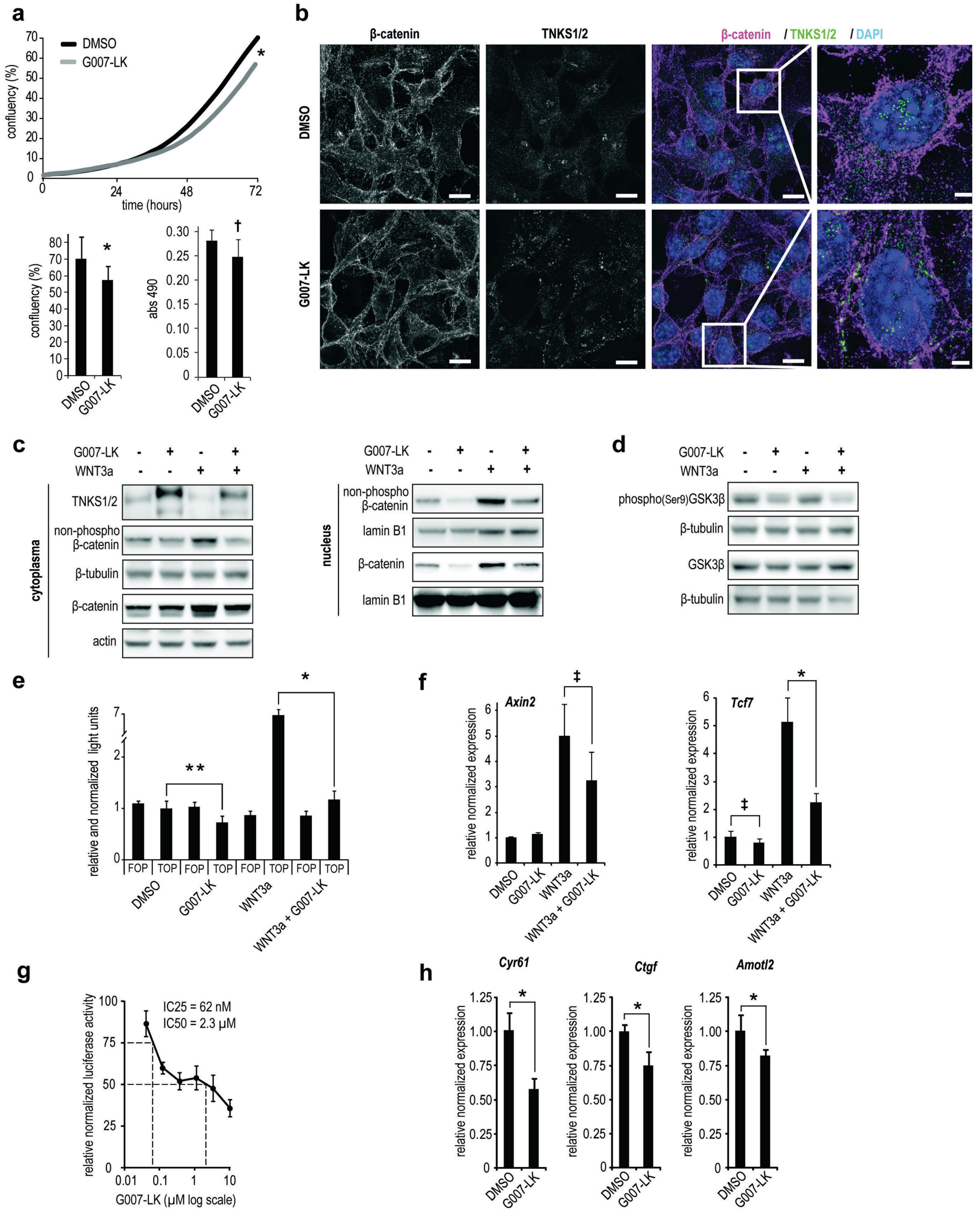

**Supplementary Fig. 1 Efficacy against biomarkers for WNT/β-catenin and YAP signaling activities upon tankyrase inhibitor treatment B16-F10 cells *in vitro*.** **a**, Real-time confluency (%) measurements of proliferating B16-F10 cells treated with vehicle (DMSO, 0.01%, black) or G007-LK (1 μM, grey). End point confluency (%, lower left panel) and MTS assay ($Abs_{490}$, lower right panel) is shown below. One-tailed t-test is indicated by * ($P < 0.01$) and Mann-Whitney rank sum test is indicated by † ($P < 0.01$). **b**, Co-staining using antibodies against β-catenin (pink) and TNKS1/2 (green) along with nuclear DAPI staining (blue) upon vehicle (0.01% DMSO) and G007-LK (1 μM) treatment (24 hours) of B16-F10 cells. The stainings show unchanged intensity of membrane β-catenin but formation of cytoplasmic TNKS1/2-containing puncta that indicates accumulation of β-catenin destruction complexes. Human-specific antibodies (mouse specific antibodies are not available) generally detect AXIN1 and AXIN2 in these puncta in human tankyrase-inhibitor treated cells. Scale bar = 10 μm. Zoom scale bar = 2 μm. **c**, Representative immunoblots of cytoplasmic TNKS1/2 as well as cytoplasmic (left) and nuclear (right) non-phospho β-catenin and total β-catenin from treated B16-F10 cell culture. Treatments used in cultured B16-F10 cells in **c**-**f**: Vehicle (DMSO, 0.01%), G007-LK (1 μM), recombinant WNT3a or WNT3a + G007-LK for 24 hours. For **c** and **d**: Actin, β-tubulin or lamin B1 document equal protein loading. **d**, Representative immunoblots of cytoplasmic and inactive form of GSK3β (phospho[ser9]GSK3β) and total GSK3β. **e**, Luciferase-based reporter assay for measuring WNT/β-catenin signaling activity. B16-F10 cells transiently transfected with superTOPflash (vector with TCF promoter binding sites) or FOPflash (control vector with mutated TCF binding sites) along with *Renilla* luciferase (for normalization). G007-LK counteracted both WNT3a-induced and non-induced superTOPflash reporter activity. One-tailed t-tests are indicated by * ($P < 0.01$) or * * ($P < 0.05$). Background SuperTOPflash versus FOPflash activities were not significantly different (two-tailed t-test, $P = 0.37$), indicating low basal WNT/β-catenin signaling activity in cultured B16-F10 cells. Mean values ± s.d. from representative experiment are shown. **f**, Real-time RT-qPCR analyses of WNT/β-catenin signaling target genes (*Axin2* and *Tcf7*). One-tailed Mann-Whitney rank sum tests are indicated by ‡ ($P < 0.05$) and one-tailed t-test is indicated by * ($P < 0.01$). Mean values ± s.d. from minimum 2 repeated measurements are shown. **g**, B16-F10 cells transiently transfected with vectors for a luciferase-based reporter assay for measuring hippo signaling activity (8XGTIIC-luciferase) and *Renilla* luciferase followed by treatment for 24 hours with vehicle (DMSO, 0.01%) or various doses of G007-LK. Pooled relative and normalized data from multiple experiments show a dose-dependent reduction in reporter activity upon G007-LK treatment. $IC_{25}$-value = 62 nM. $IC_{50}$-value = 2.3 μM. **h**, Real-time RT-qPCR analyses of YAP signaling target genes (*Cyr61*, *Ctgf* and *Amotl2*) from B16-F10 cell culture treated (24 hours) with vehicle (DMSO, 0.01%) or G007-LK (1 μM). One-tailed Mann-Whitney rank sum tests are indicated by † ($P < 0.01$) and one-tailed t-tests are indicated by * ($P < 0.01$). Mean values ± s.d. from minimum 2 repeated measurements are shown.

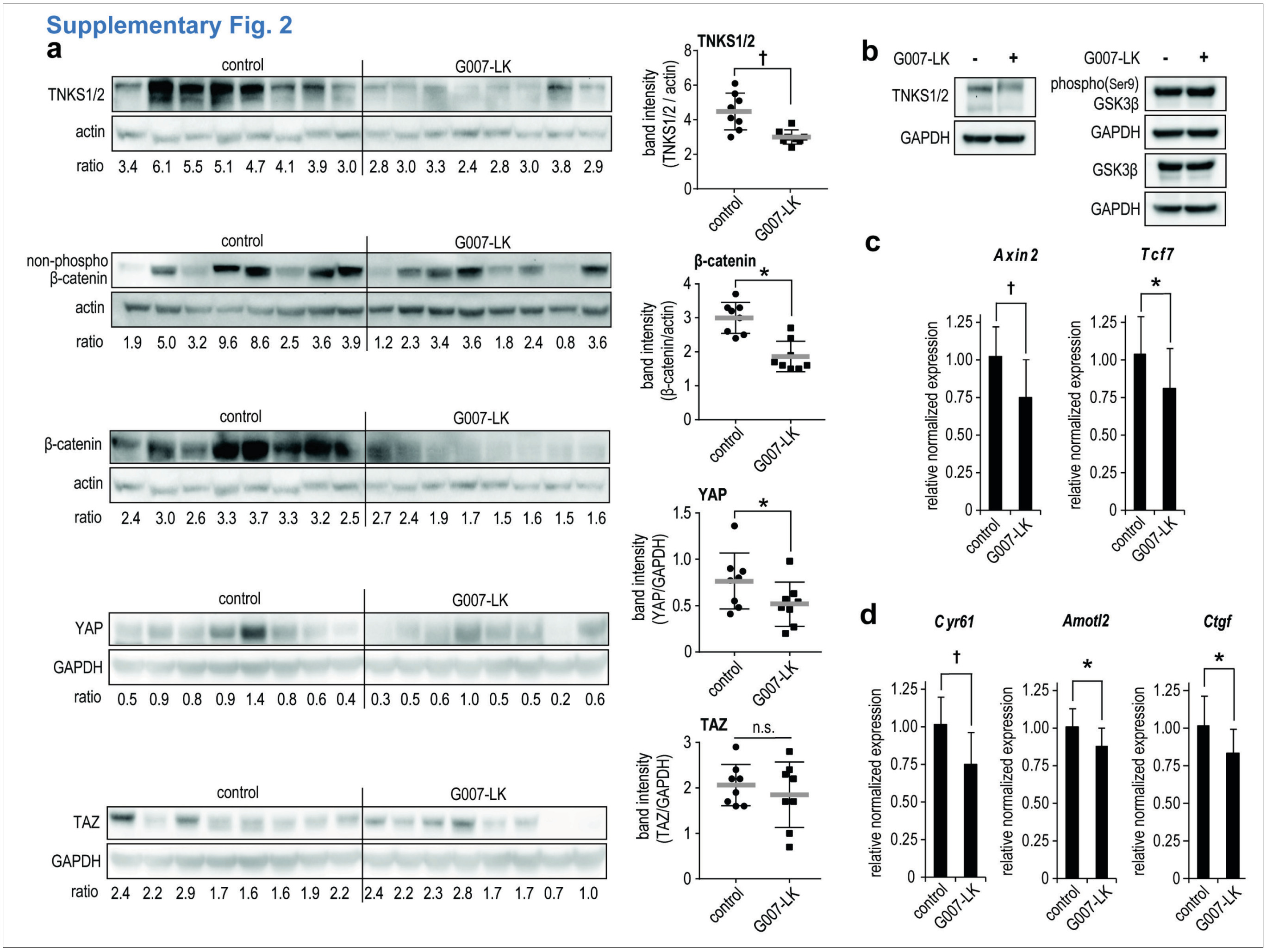


**Supplementary Fig. 2 Efficacy against biomarkers for WNT/β-catenin and YAP signaling activities upon tankyrase inhibitor treatment of B16-F10 tumors in C57BL/6N mice. a**, Representative immunoblots from whole s.c. B16-F10 tumors showing reduced expression of TNKS1/2 levels, active β-catenin (non-phospho, Ser33/37/Thr41, mid panel), β-catenin (total), YAP and TAZ upon 4 days of treatment with G007-LK diet compared to controls (both groups, $n$ = 8). Quantified ratios (protein vs. actin loading control) are shown below the graphics and plotted in the right panels for TNKS1/2, β-catenin, YAP and TAZ. One-tailed Mann-Whitney rank sum test is indicated by † ($P < 0.01$) and one-tailed t-tests are indicated by * ($P < 0.01$). n.s. = not statistically significant. **b**, Pooled ($n$ = 8) samples from G007-LK-treated whole B16-F10 tumors probed against TNKS1/2, phospho(ser9)GSK3β and GSK3β. GAPDH documents equal protein loading. **c**, Real-time RT-qPCR analyses of WNT/β-catenin signaling target genes (*Axin2* and *Tcf7*). For **c** and **d**: From G007-LK-treated s.c. B16-F10 tumors versus controls ($n$ = 8). One-tailed Mann-Whitney rank sum tests are indicated by † ($P < 0.01$) and one-tailed t-tests are indicated by * ($P < 0.01$). Mean values ± s.d. for 2 repeated measurements are shown. **d**, Real-time RT-qPCR analyses of YAP signaling target genes (*Cyr61*, *Amotl2* and *Ctgf*).

#### Supplementary Fig. 3

##### B16F-10 tumors in in C57BL/6N mice

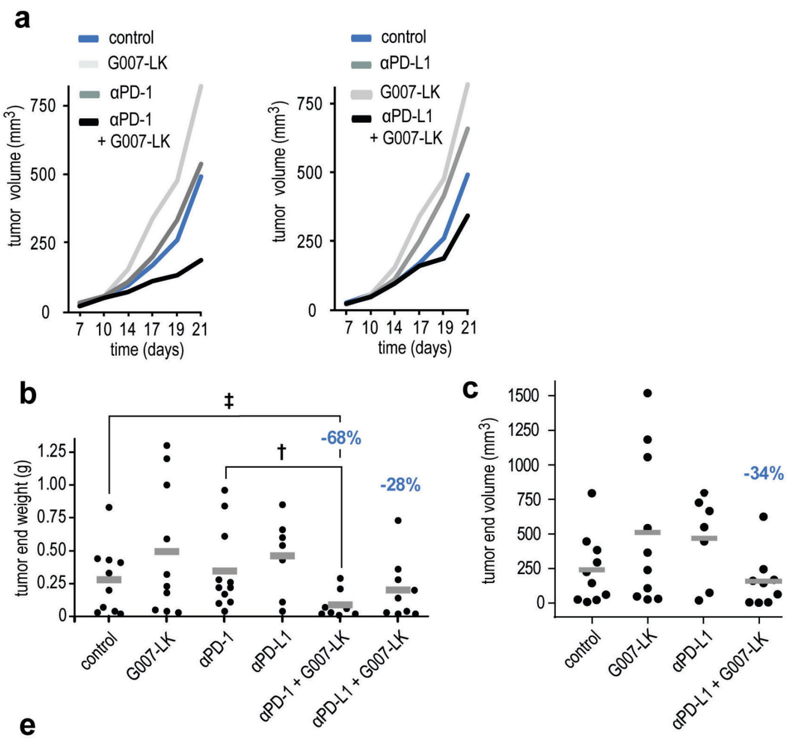


d

| | mean mm3 | mean mm3 (%) | Δ mean mm3 (%) | mean g | mean g (%) | Δ mean g (%) |
|---|---|---|---|---|---|---|
| ctrl | 240 | 100 | 0 | 0.3 | 100 | 0 |
| G007-LK | 512 | 213 | 113 | 0.5 | 176 | 76 |
| αPD-1 | 364 | 151 | 51 | 0.4 | 124 | 24 |
| αPD-L1 | 468 | 195 | 95 | 0 | 165 | 65 |
| αPD-1 + G007-LK | 40 | 17 | -83 | 0.1 | 32 | -68 |
| αPD-L1 + G007-LK | 158 | 66 | -34 | 0.2 | 72 | -28 |

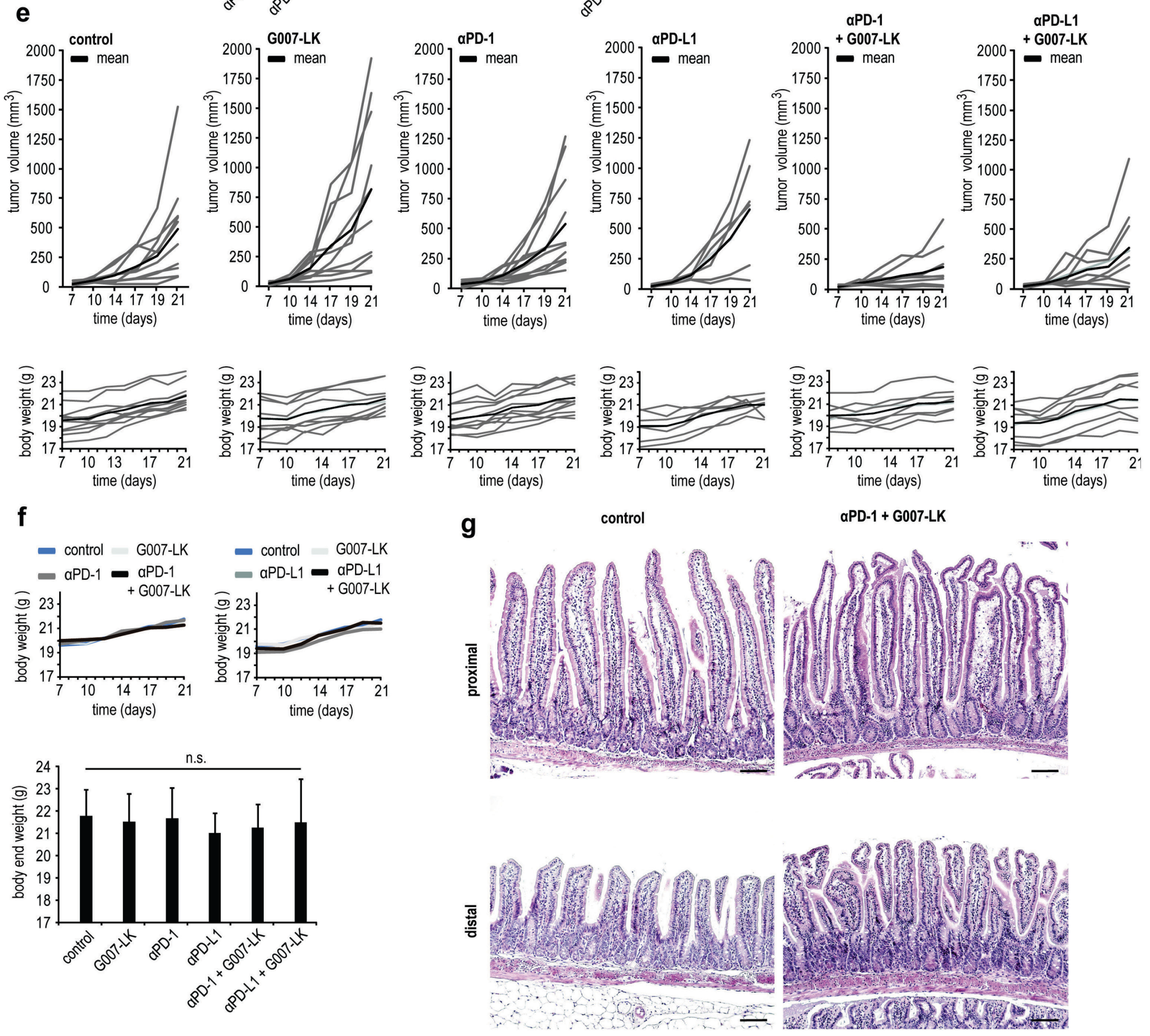

**Supplementary Fig. 3 Anti-tumor efficacy of combined tankyrase and checkpoint inhibitor treatment of B16-F10 tumors in C57BL/6N mice. a**, Mean s.c. B16-F10 tumor volumes ($mm^3$). For **a** and **f**: Control diet (blue), G007-LK diet (light grey, left panel), anti-PD-1 (grey, left panel) and anti-PD-1/G007-LK (black, left panel), anti-PD-L1 (grey, right panel) and anti-PD-L1/G007-LK (black, right panel). For **a**-**f**: Mice treated from day 10 until day 21. Treatments: Control diet ($n$ = 10), G007-LK diet ($n$ = 10), anti-PD-1 ($n$ = 11), anti-PD-L1 ($n$ = 7), anti-PD-1/G007-LK ($n$ = 8) and anti-PD-L1/G007-LK ($n$ = 9). **b**, Tumor end weight reduction upon anti-PD-1/G007-LK (-68%) or anti-PD-L1/G007-LK (-28%) treatment compared to control. Mann-Whitney rank sum tests are indicated by $^{‡}$ ($P$ < 0.05) and $^{†}$ ($P$ < 0.01). **c**, Tumor end volume reduction upon anti-PD-L1/G007-LK (-34%) treatment compared to control. **d**, Left table depicts mean tumor end volumes ($mm^3$), mean relative tumor end volumes ($mm^3$ [%]) and relative differences when compared to control ($\Delta mm^3$ [%]). Right table depicts mean tumor end weights (g), mean relative tumor end weights (g [%]) and relative differences when compared to control (Δg [%]). **e**, Single s.c. B16-F10 tumor volumes (upper panels) and body weights (lower panels). Mean values are shown in black. **f**, No differences in mean body weights in mice treated from day 10 until day 21 (upper panels) and body end weight (lower panel). n.s = not significant. **g**, A histopathological examination performed at experiment termination documented no abnormalities in morphology, proliferation or differentiation in the small intestinal mucosa upon combined anti-PD-1/G007-LK treatment. Representative pictures from multiple sections of H&E-stained proximal (upper panels) and distal (lower panels) parts of the small intestine from mice treated with control diet (left panels, $n$ = 6) or anti-PD-1/G007-LK treatment (right panels, $n$ = 6). Scale bars: 100 µm (original magnification ×100).

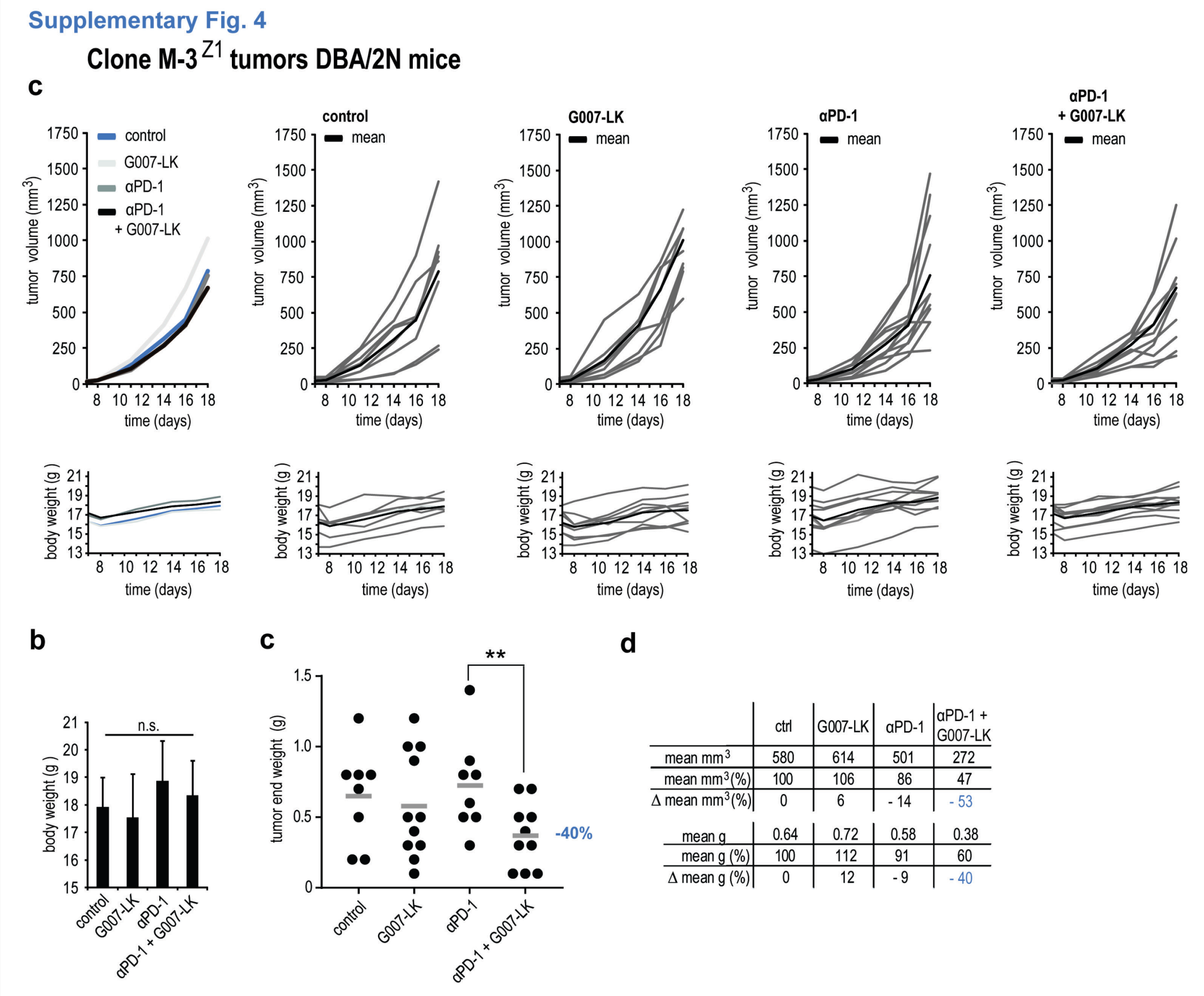


| | ctrl | G007-LK | αPD-1 | αPD-1 + G007-LK |
|---|---|---|---|---|
| mean mm³ | 580 | 614 | 501 | 272 |
| mean mm³(%) | 100 | 106 | 86 | 47 |
| Δ mean mm³(%) | 0 | 6 | - 14 | - 53 |
| mean g | 0.64 | 0.72 | 0.58 | 0.38 |
| mean g (%) | 100 | 112 | 91 | 60 |
| Δ mean g (%) | 0 | 12 | - 9 | - 40 |

**Supplementary Fig. 4 Anti-tumor efficacy of combined tankyrase and checkpoint inhibitor treatment of Clone M-3$^{Z1}$ tumors DBA/2N mice. a**, Clone M-3$^{Z1}$ mean (leftmost panels) and single (right panels) tumor (s.c.) volumes (upper panels) and body weights (lower panels) from mice treated from day 8 until day 18 with control diet (*n* = 8, in blue), G007-LK diet (*n* = 9, in light grey), anti-PD-1 (*n* = 11, in dark grey) and combinations of anti-PD-1/G007-LK (*n* = 11, in black). Mean values are indicated by black lines. **b**, No statistically significant (n.s.) differences in mean body end weight. **c**, Clone M-3$^{Z1}$ tumor end weight reduction upon anti-PD-1/G007-LK treatment (-40%) compared to control (two-tailed t-test = 0.085). Two-tailed t-test is indicated by ** ($P < 0.05$). Mean values are indicated by grey lines ± s.d.. **d**, Upper table depicts mean tumor end volumes (mm$^3$), mean relative tumor end volumes (mm$^3$ [%]) and relative differences when compared to control (Δmm$^3$ [%]). Lower table depicts mean tumor end weights (g), mean relative tumor end weights (g [%]) and relative differences when compared to control (Δg [%]).

#### Supplementary Fig. 5

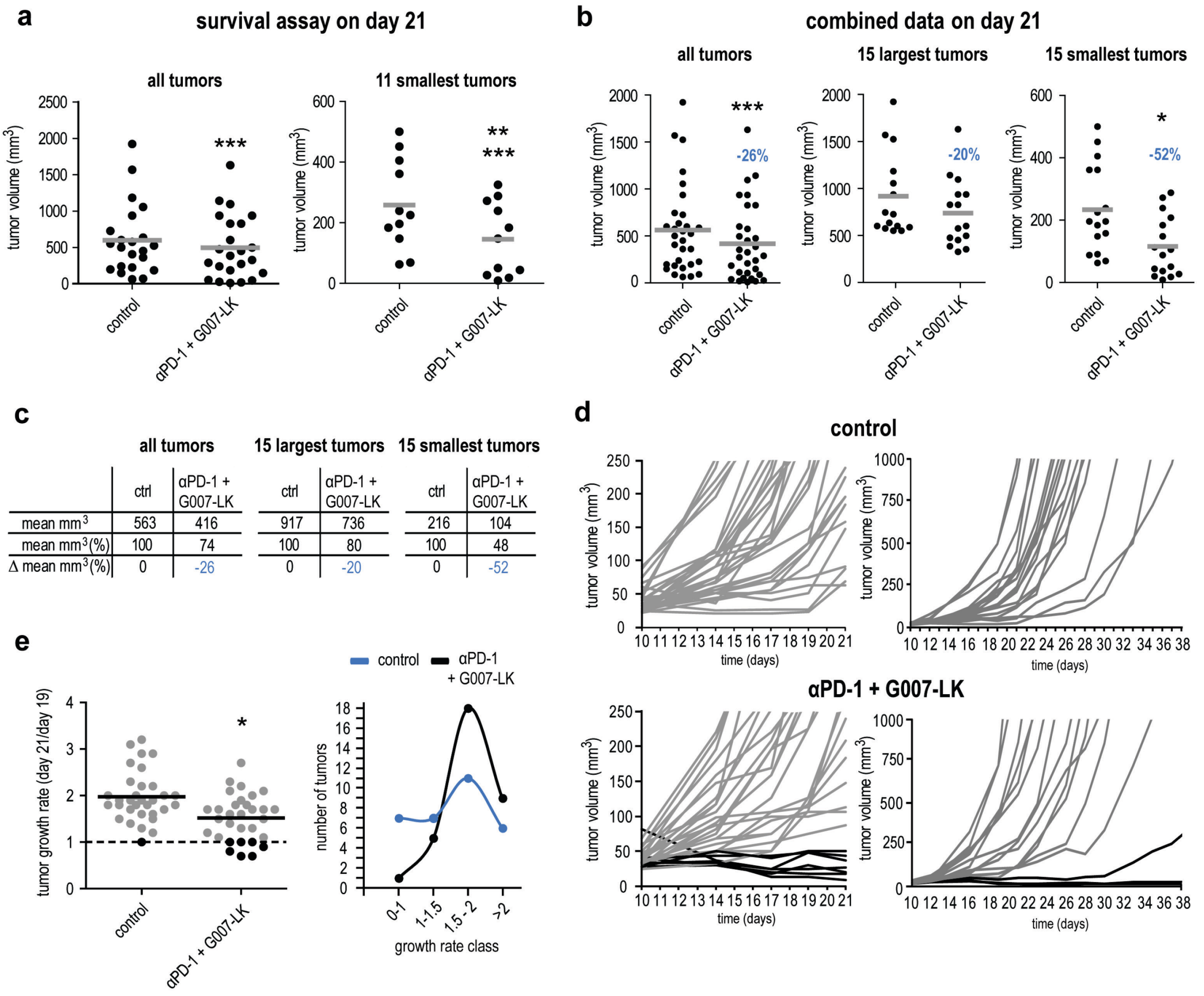

**Supplementary Fig. 5 Anti-tumor efficacy of combined tankyrase and checkpoint inhibitor treatment of B16-F10 tumors in C57BL/6N mice in a survival assay. a**, B16-F10 tumor (s.c.) volume reduction for the survival assay on day 21 upon combined anti-PD-1/G007-LK treatment ($n$ = 23) compared to control ($n$ = 22). For all tumors (left panel): Paired t-test is indicated by *** ($P$ < 0.001). For the 11 smallest tumors in each group (right panel): Paired t-test is indicated by *** ($P$ < 0.001) and one-tailed t-test is indicated by ** ($P$ < 0.05). Mean values are indicated by grey lines ± s.d.. **b**, Plots showing combined data for tumor volume reduction from the tumor growth and survival assays on day 21 upon combined anti-PD-1/G007-LK treatment ($n$ = 32) compared to control ($n$ = 31). For all tumors (left panel, 26% reduction, paired t-test is indicated by *** [$P$ < 0.001]), the 15 largest tumors in each group (20% reduction, mid panel) and the 15 smallest tumors in each group (52% reduction, one-tailed t-test is indicated by * [$P$ < 0.01]). **c**, Tables depicting mean tumor end volumes ($mm^3$), relative mean tumor end volumes ($mm^3$ [%]) and relative differences when compared to control ($\Delta mm^3$ [%]) for all tumors (left panel) along with the 15 largest (mid panel) and smallest (right panel) tumors in each group. **d**, Graphs showing caliper-based measurements of single s.c. tumor volumes from mice treated from day 10 until day 21 in the survival assay. Control (upper panels) and combined anti-PD-1/G007-LK treatment (lower panels). Left panels (Y-axis, 0-250 $mm^3$), black lines depict tumors with volumes scoring below 50 $mm^3$ (7 out of 31 = 22.5%). Right panels (Y-axis, 0-1000 $mm^3$) with black lines depicting tumors from animals alive on day 38 (3 out of 16 = 18.75%). **e**, Decrease in tumor growth rate (day 21/day19 = growth rate) for combined anti-PD-1/G007-LK treatment (growth rate = 1.5, $n$ = 23) compared to control (growth rate = 2.0, $n$ = 22). Plot in left panel: One-tailed t-test is indicated by * ($P$ < 0.01) and black color indicates tumors with growth rates ≤ 1. Graphs in right panel: Number of tumors in each growth rate class: 0-1, 1-1.5, 1.5-2 and >2. Control in blue and anti-PD-1/G007-LK treatment in black.

#### Supplementary Fig. 6

**a**

##### *Ctnnb1* exon 4

**Coding sequence**
ATATTGACGGGCAGTATGCAATGACTAGGGCTCAGAGGGTCCGAGCTGCCATGTTCCCT
GAGACGCTAGATGAGGGCATGCAGATCCCATCCACGCAGTTTGACGCTGCTCATCCCAC
TAATGTCCAGCGCTTGGCTGAACCATCACAGATGTTGAAACATGCAGTTGTCAATTTGAT
TAACTATCAGGATGACGCGGAACTTGCCACACGTGCAATTCCTGAGCTGACAAAACTGC
TAAACGATGAGGACCAG

**Amino acid sequence**
IDGQYAMTRAQRVRAAMFPETLDEGMQIPSTQFDAAHPTNVQRLAEPSQMLKHAVVNLINY
QD**DAELATRAIPELTKLLNDEDQ**

---

##### *Ctnnb1* KO1

**Allele #1,** single base loss > frameshift and premature stop:
TATCAGGAT-ACGCGGAACTTGCCACACGTGCAATTCCT
**Amino acid sequence**
IDGQYAMTRAQRVRAAMFPETLDEGMQIPSTQFDAAHPTNVQRLAEPSQMLKHAVVNLINY
QD**TRNLPHVQFLS***

**Allele #2,** Insertion G - frameshift and premature stop:
TCAATTTGATTAACTATCAGGATggACGCGGA
**Amino acid sequence**
IDGQYAMTRAQRVRAAMFPETLDEGMQIPSTQFDAAHPTNVQRLAEPSQMLKHAVVNLINY
QD**GRGTCHTCNS***

---

##### *Ctnnb1* KO2

**Allele #1**
tg > c- > frameshift and premature stop:
TCAGGAc-ACGCGGAACTTGC
**Amino acid sequence**
IDGQYAMTRAQRVRAAMFPETLDEGMQIPSTQFDAAHPTNVQRLAEPSQMLKHAVVNLINY
QD**TRNLPHVQFLS***

**Allele #2**
tg > a- frameshift and premature stop:
TAACTATCAGGAA-AC
**Amino acid sequence**
IDGQYAMTRAQRVRAAMFPETLDEGMQIPSTQFDAAHPTNVQRLAEPSQMLKHAVVNLINY
Q**ETRNLPHVQFLS***

**b**

**c**

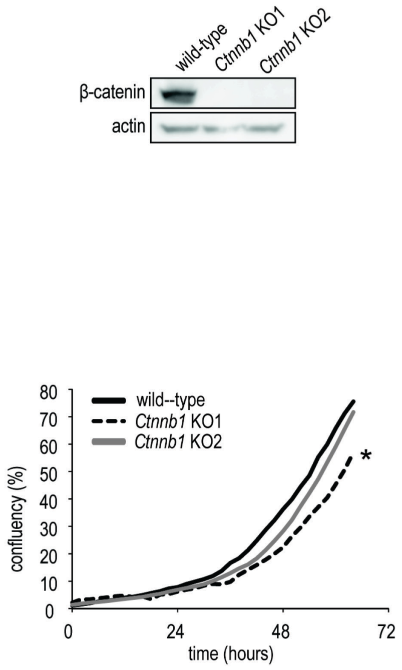


**d**

**e**

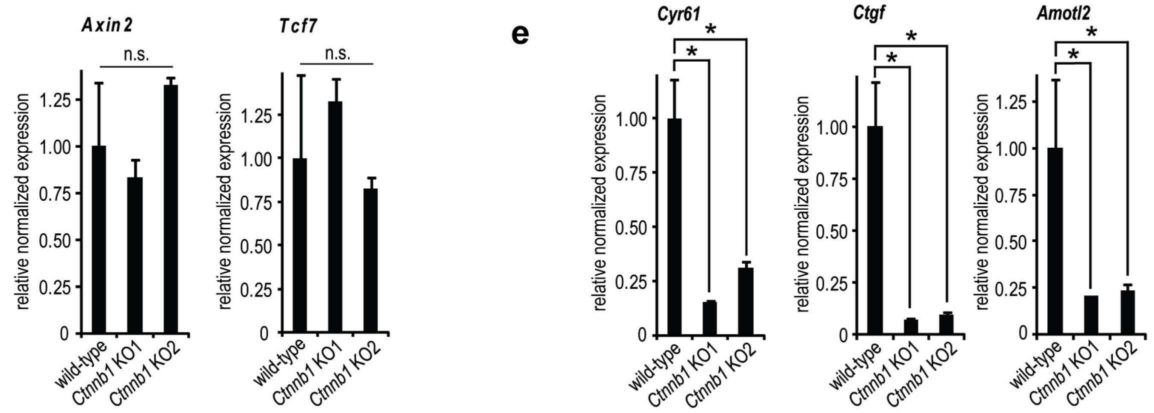

**Supplementary Fig. 6 CRISPR/Cas9-based knock-out of Ctnnb1 in B16-F10 cells. a**, Coding (upper) and amino acid (lower) sequences for wild-type *Ctnnb1* exon 4. Alterations in coding and amino acid (highlighted in blue) sequences in allele 1 and 2 upon CRISPR-Cas9-based gene editing are shown for B16-F10$^{Ctnnb1\mathrm{KO}}$ clone 1 and 2 (*Ctnnb1* KO1 and 2). **b**, Immunoblots from total lysates showing loss of β-catenin in B16-F10$^{Ctnnb1\mathrm{KO}}$ cell lines compared to wild-type B16-F10 cells. Actin is used as loading control. **c**, Measurements of real-time confluence (%) for proliferating B16-F10 wild-type and B16-F10$^{Ctnnb1\mathrm{KO}}$ cell lines. *Ctnnb1* KO1 displays moderately reduced proliferation speed compared to wild-type B16-F10 cells. Two-tailed t-test is indicated by * ($P < 0.01$). **d**, Real-time RT-qPCR analyses of B16-F10$^{Ctnnb1\mathrm{KO}}$ cell lines show no regulation of WNT/β-catenin signaling target gene expression (*Axin2* and *Tcf7*) compared to wild-type B16-F10 cells. Mean values ± s.d. from minimum 2 repeated measurements are shown and n.s. = not significant. Low basal WNT/β-catenin signaling activity in cultured B16-F10 cells (see Supplementary Fig. 1e and f) could not be further reduced in B16-F10$^{Ctnnb1\mathrm{KO}}$ cell lines when compared to wild-type B16-F10 cells. **e**, Real-time RT-qPCR analyses of B16-F10$^{Ctnnb1\mathrm{KO}}$ cell lines show down-regulated YAP signaling target gene expression (*Cyr61*, *Ctgf* and *Amotl2*) compared to wild-type B16-F10 cells. Two-tailed t-tests are indicated by * ($P < 0.01$). Mean values ± s.d. from minimum 2 repeated measurements are shown.

**Supplementary Fig. 7**

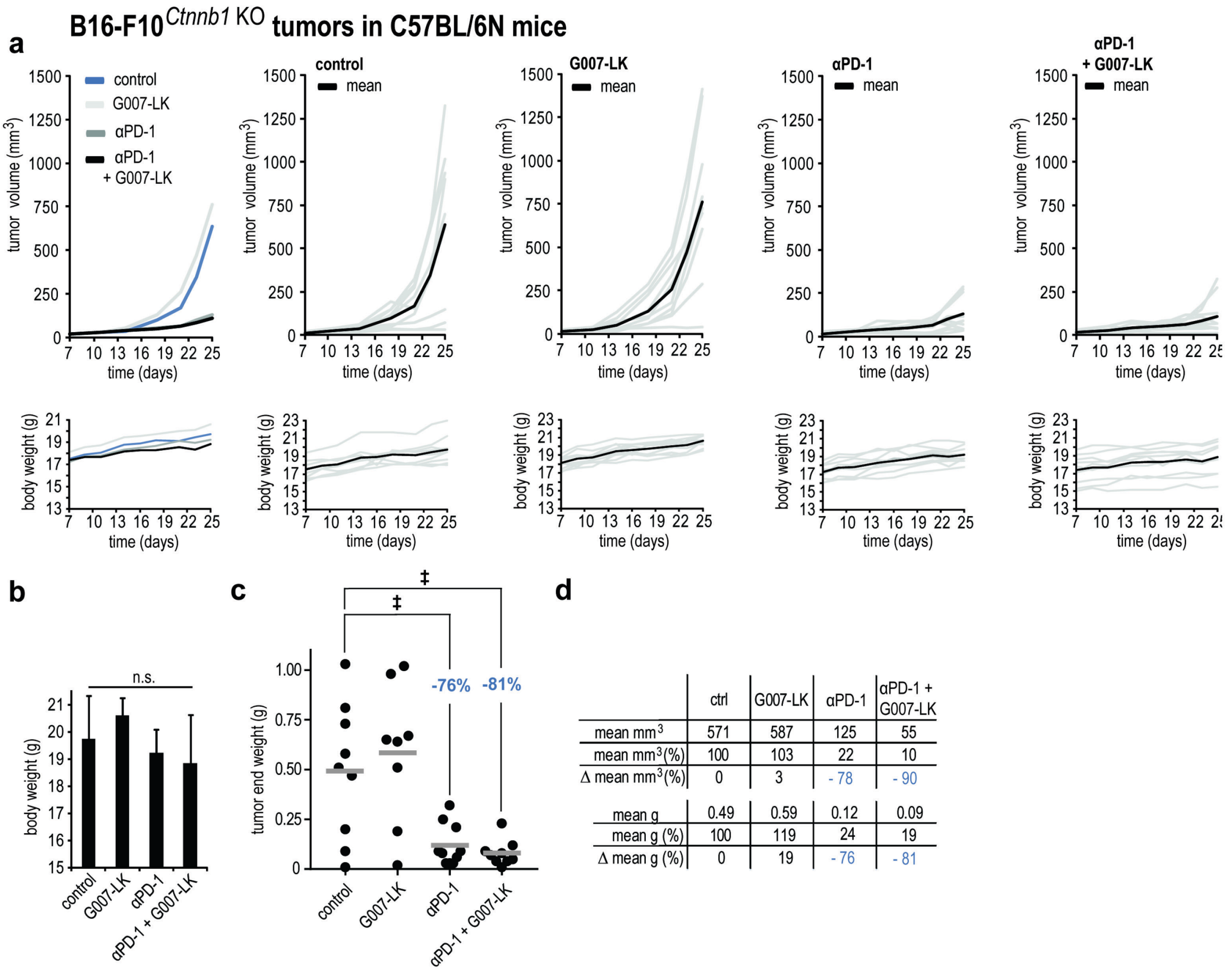


| | ctrl | G007-LK | αPD-1 | αPD-1 + G007-LK |
|---|---|---|---|---|
| mean mm$^3$ | 571 | 587 | 125 | 55 |
| mean mm$^3$(%) | 100 | 103 | 22 | 10 |
| Δ mean mm$^3$(%) | 0 | 3 | - 78 | - 90 |
| mean g | 0.49 | 0.59 | 0.12 | 0.09 |
| mean g (%) | 100 | 119 | 24 | 19 |
| Δ mean g (%) | 0 | 19 | - 76 | - 81 |

**Supplementary Fig. 7 B16-F10*Ctnnb1*KO tumors are sensitive to PD-1 inhibition in C57BL/6N mice. a**, B16-F10*Ctnnb1*KO mean (leftmost panels) and single (right panels) tumor (s.c.) volumes (upper panels) and body weights (lower panels) from mice treated from day 11 until day 25 with control diet ($n$ = 9, in blue), G007-LK diet ($n$ = 9, in light grey), anti-PD-1 ($n$ = 10, in dark grey) and combinations of anti-PD-1/G007-LK ($n$ = 10, in black). Mean values are indicated by black lines. **b**, Graph showing no statistically significant (n.s.) differences in mean body end weight. **c**, B16-F10*Ctnnb1*KO tumor (s.c.) end weight reduction upon anti-PD-1 (-76%) and combined anti-PD-1/G007-LK treatment (-81%). Mann-Whitney rank sum tests are indicated by ‡ ($P$ < 0.05). Mean values are indicated by grey lines ± s.d.. **d**, Upper table depicts mean tumor end volumes (mm$^3$), mean relative tumor end volumes (mm$^3$ [%]) and relative differences when compared to control (Δmm$^3$ [%]). Lower table depicts mean tumor end weights (g), mean relative tumor end weights (g [%]) and relative differences when compared to control (Δg [%]).

#### Supplementary Fig. 8

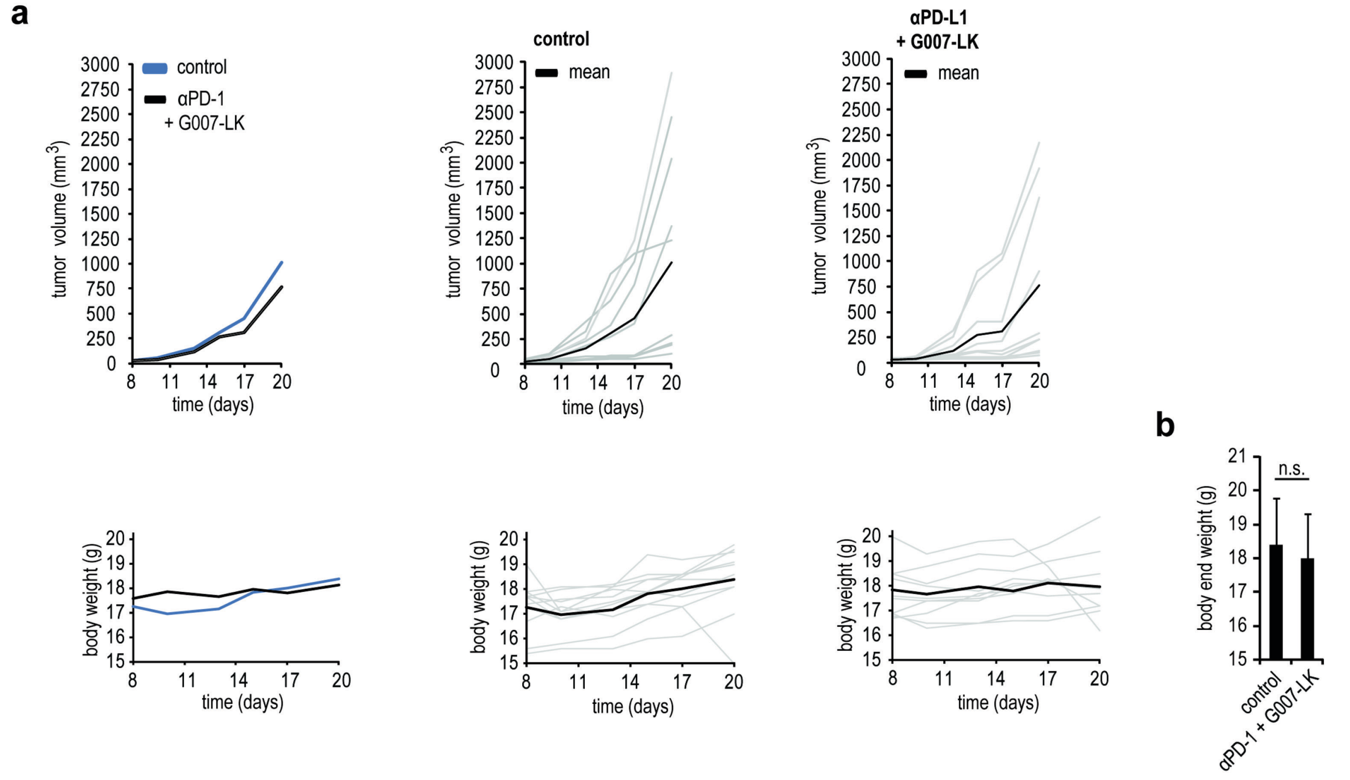


**Supplementary Fig. 8 No anti-tumor efficacy of combined tankyrase and checkpoint inhibitor treatment of B16-F10 tumors in Rag2$^{-/-}$ mice. a**, Mean B16-F10 (s.c. in Rag2$^{-/-}$ mice )(leftmost panels) and single (right panels) tumor volumes (upper panels) and in addition body weights (lower panels) from mice treated from day 8 until day 20 with control diet (*n* = 11, in blue) or anti-PD-1/G007-LK (*n* = 10, in black). Mean values are indicated by black lines (right panels). **c,** Graph showing no statistically significant difference in mean body end weight. n.s. = not significant.

#### Supplementary Figure 9

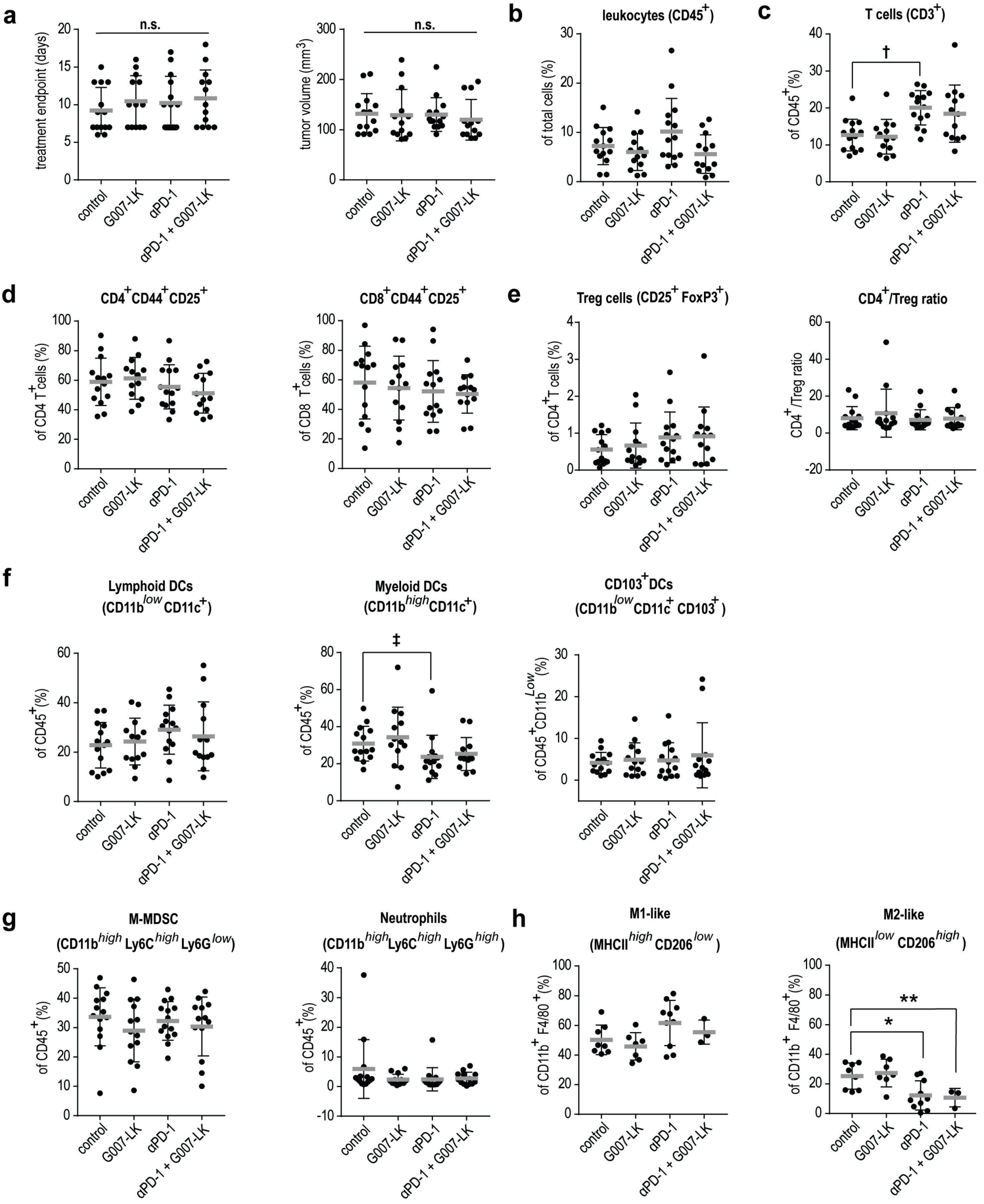

**Supplementary Fig. 9 Anti-PD-1 treatment induces quantitative impact on intratumoral leukocyte and lymphocyte subsets in B16-F10 tumors.** **a**, Collection time-points (after 7-17 days of treatment, left panel) and tumor volumes (right panel) for s.c. tumors isolated for flow cytometry analysis (shown in **b**-**g**) from B16-F10-challenged C57BL/6N mice treated with control diet (*n* =14), G007-LK diet (*n* = 13), anti-PD-1 (*n* = 14) or anti-PD-1/G007-LK (*n* = 13). n.s. = not significant. For **a**-**h**, mean values are indicated by grey lines ± s.d.. **b**, Total leukocytes ($CD45^+$) shown as % of total cell number. **c**, T cells ($CD3^+$) shown as % of leukocytes ($CD45^+$). **d**, Quantitation of T-cell subsets: $CD4^+CD44^+CD25^+$ T-cells shown as % of $CD4^+$ T-cells, $CD8^+CD44^+CD25^+$ T-cells shown as % of $CD8^+$ T-cells. **e**, $T_{reg}$ ($CD25^+FoxP3^+$) cells plotted as % of $CD4^+$ T-cells and ratio of $CD4^+$ T-cells to $T_{reg}$ ($CD25^+FoxP3^+$) cells, (CD4+/$T_{reg}$ ratios). **f**, Lymphoid DCs ($CD45^+CD11b^{low}CD11c^+$) and myeloid DCs ($CD45^+CD11b^{high}CD11c^+$) shown as % of leukocytes ($CD45^+$). $CD103^+$DCs ($CD45^+CD11b^{low}CD11c^+CD103^+$) shown as % of $CD11b^{low}$ leukocytes. Mann-Whitney rank sum test is indicated by $^‡$ ($P < 0.05$). **g**, Quantitation of sub-populations of myeloid-derived suppressor cells: M-MDSCs ($CD11b^{high}Ly6C^{high}Ly6G^{low}$) and neutrophils ($CD11b^{high}Ly6C^{high}Ly6G^{high}$) shown as % of leukocytes ($CD45^+$). **h**, M1-like macrophages ($MCHII^{high}CD206^{low}$) and M2-like macrophages ($MCHII^{low}CD206^{high}$) as % of $CD11b^{high}F4/80^+$ macrophages from single-cell suspension of s.c. tumors isolated after 11 days of therapy with control diet (*n* = 8), G007-LK diet (*n* = 7), anti-PD-1 (*n* = 10) or anti-PD-1/G007-LK (*n* = 3) in B16-F10-challenged C57BL/6N mice. One way ANOVA (Holm-Sidak method versus control) indicated by * ($P < 0.05$), two-tailed t-test is indicated by ** ($P < 0.05$).

#### Supplementary Fig. 10

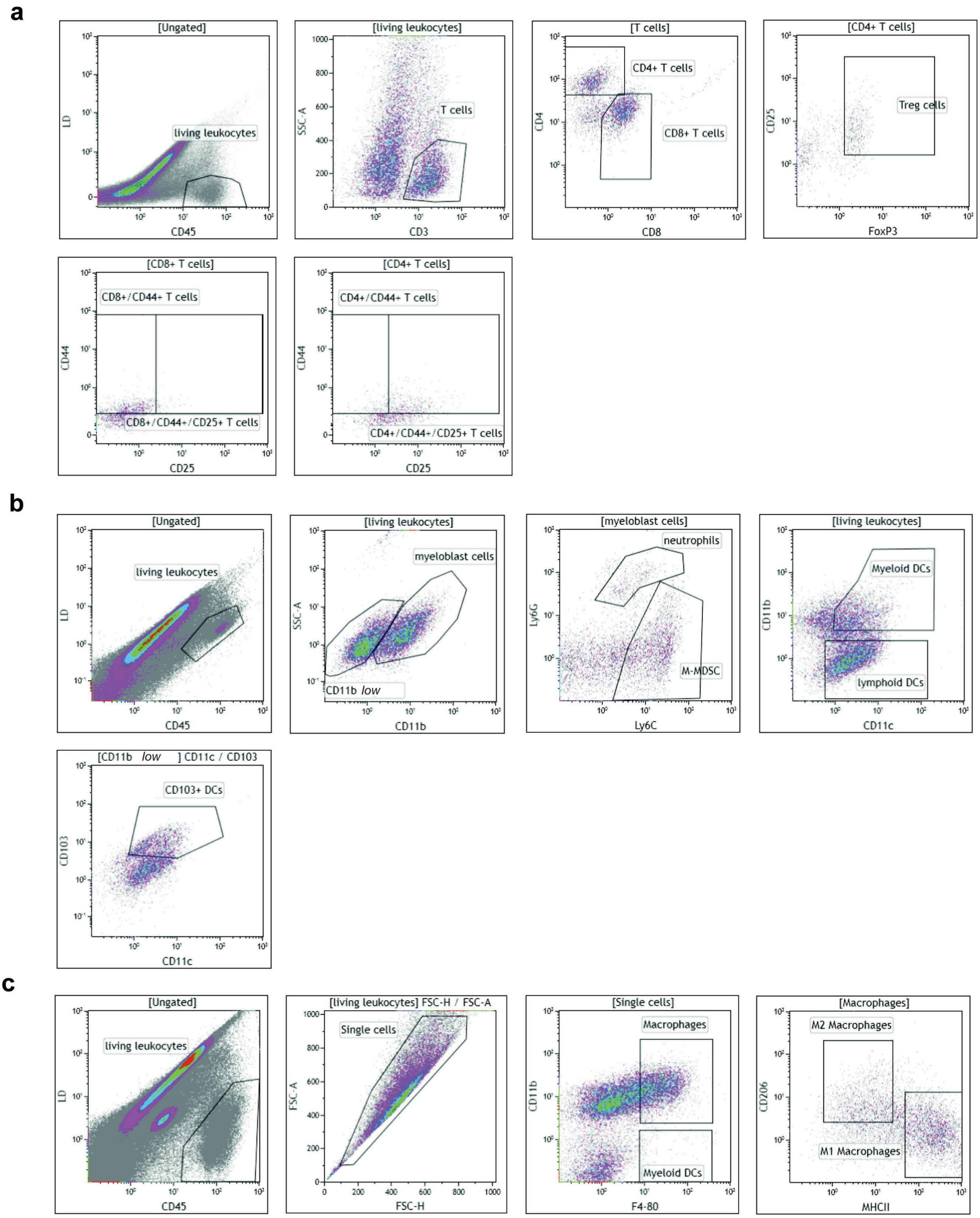

**Supplementary Fig. 10 Flow cytometry gating strategy for quantification of intratumoral leukocyte and lymphocyte subsets in treated B16-F10 tumors.** Representative examples of flow cytometry gating strategy using primary material from s.c. B16-F10 tumors treated with DMSO, G007-LK, anti-PD-1 or anti-PD-1/G007-LK for determining sub-populations of T cells, myeloid-derived suppressor cells and macrophages (see Fig. 3 and Supplementary Fig. 9)**.** The gating strategies were defined by ProQinase[49,50] (Germany). **a**, Gating strategy for T cells: Living leukocytes (CD45$^{+}$), T cells (CD3$^{+}$), CD4$^{+}$ T cells (CD3$^{+}$CD4$^{+}$), CD8+ T cells (CD3$^{+}$CD8$^{+}$), Treg cells (CD3$^{+}$CD4$^{+}$CD25$^{+}$FoxP3$^{+}$), activated CD4$^{+}$ T cells (CD4$^{+}$CD44$^{+}$CD25$^{+}$) and activated CD8$^{+}$ T cells (CD8$^{+}$CD44$^{+}$CD25$^{+}$). **b**, Gating strategy for myeloid-derived suppressor cells: Living leukocytes (CD45$^{+}$), M-MDSCs (CD45$^{+}$CD11b$^{high}$Ly6C$^{\textit{high}}$Ly6G$^{\textit{low}}$), neutrophils (CD45$^{+}$CD11b$^{high}$Ly6C$^{\textit{high}}$Ly6C$^{\textit{high}}$), myeloid DCs (CD45$^{+}$CD11b$^{high}$CD11c$^{+}$), lymphoid DCs (CD45$^{+}$CD11b$^{low}$CD11c$^{+}$), CD103$^{+}$ DCs (CD45$^{+}$CD11b$^{\textit{low}}$CD11c$^{+}$CD103$^{+}$). **c**, Gating strategy for macrophages: Living leukocytes (CD45$^{+}$), macrophages (CD45$^{+}$CD11b$^{high}$F4/80$^{+}$), M1-like macrophages (CD45$^{+}$CD11b$^{high}$F4/80$^{+}$MHCII$^{\textit{high}}$CD206$^{\textit{low}}$) and M2-like macrophages (CD45$^{+}$CD11b$^{high}$F4/80$^{+}$MHCII$^{\textit{low}}$CD206$^{\textit{high}}$).

#### Supplementary Fig. 11

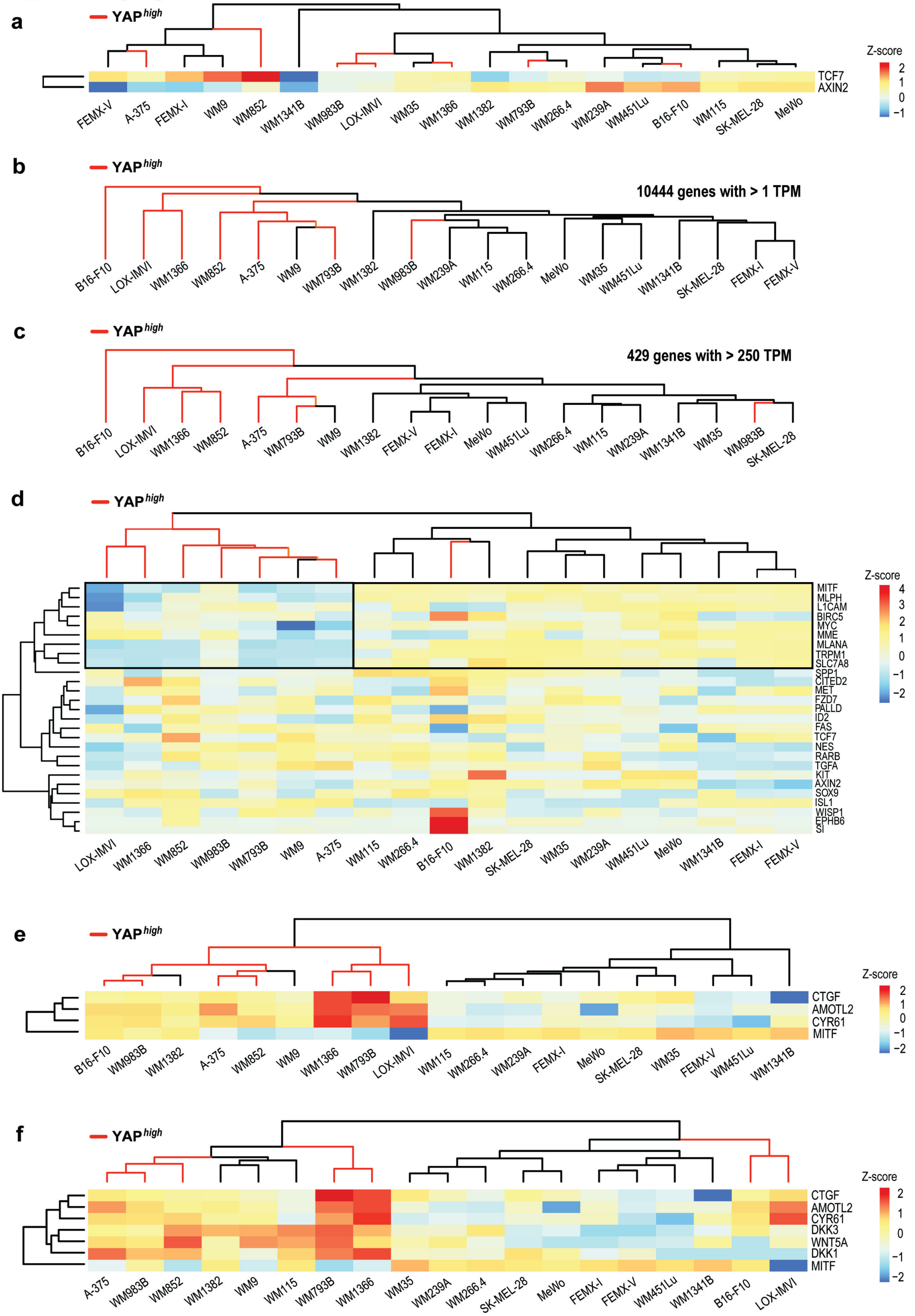

**Supplementary Fig. 11 High relative YAP signaling activity correlates with low baseline *MITF* expression.** **a**, Heatmap and clustering of transcribed WNT/β-catenin signaling target genes (*Axin2* and *Tcf7*) for 18 untreated human melanoma cell lines (pooled triplicates used for RNA sequencing) along with murine B16-F10 melanoma (biological triplicates used for RNA sequencing). For **a**-**f**: Scale bar indicates relative differences in Z-score values for log2 TPMs within each row, all values are from untreated samples. Samples displaying high relative transcription of YAP signaling target genes ($YAP^{high}$, see Fig 4a) are highlighted by orange branches in the dendrogram. **b**, Clustering of 10444 transcribed genes with >1 TPMs. **c**, Clustering of 429 transcribed genes with >250 TPMs. Murine B16-F10 does not display cross-species clustering with human samples. **d**, Heatmap and clustering of a panel of 27 markers for β-catenin-controlled melanoma cell fate and proliferation[24]. 6 of 7 samples in the $YAP^{high}$ subset cluster together, - and they cluster particularly together caused by relative lower expression of *MITF* as well as *MLPH*, *MLANA*, *TRPM1* and *SLC7A8* (highlighted by black box). **e**, Heatmap and clustering of transcribed YAP signaling target genes (*Cyr61*, *Ctgf* and *Amotl2*) versus *MITF*. $YAP^{high}$ correlates with relative low *MITF* transcription ($MITF^{low}$). **f**, Heatmap and clustering showing that high relative levels of YAP signaling activity coincides with high expression of DKK3, WNT5a and DKK1 as well as low *MITF* transcription.

#### Supplementary Fig. 12

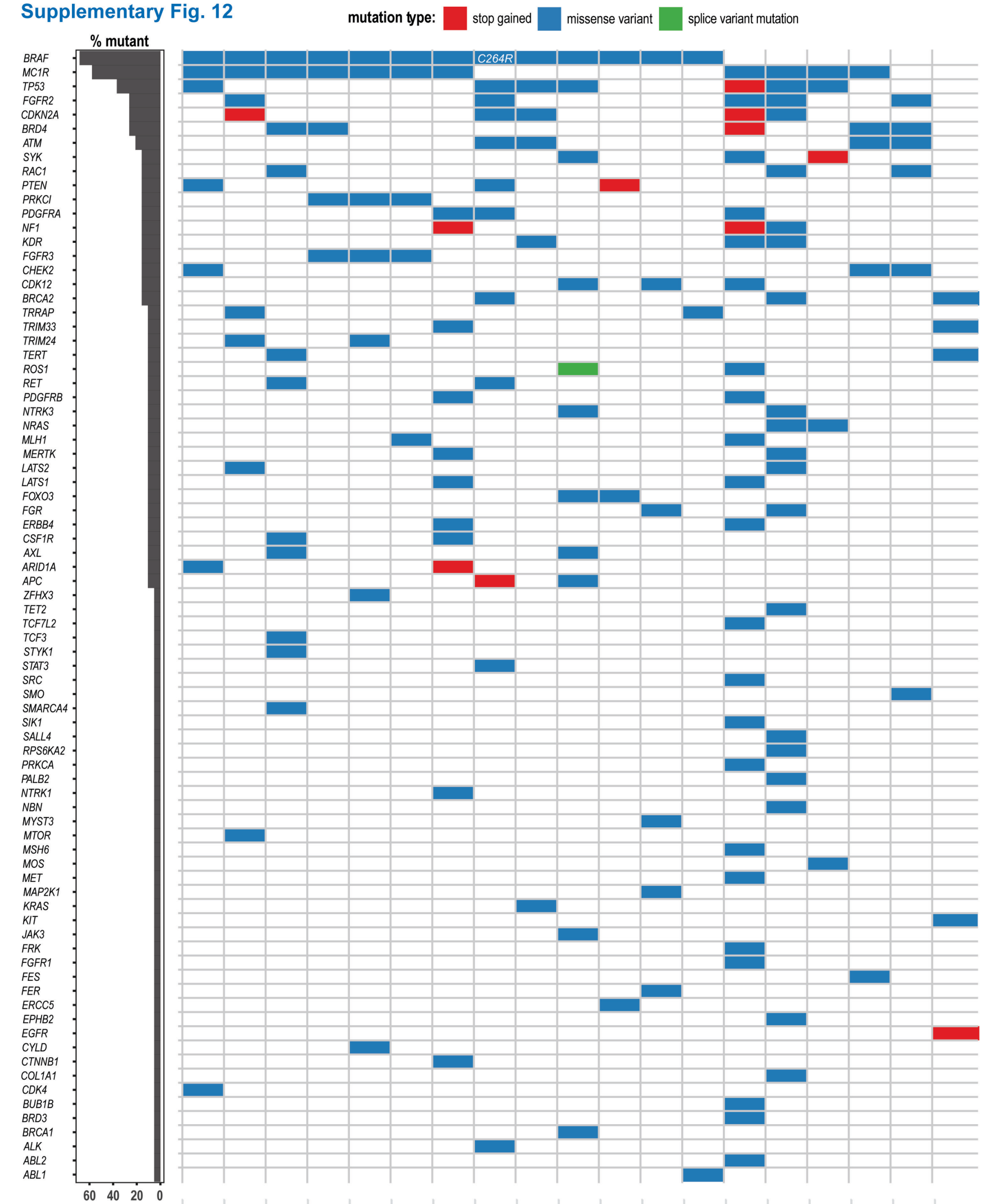


| | SK-MEL-28 | A-375 | WM9 | WM239A | WM266.4 | WM115 | LOX-IMVI | B16-F10 | WM983B | WM451Lu | WM793B | WM1341B | WM35 | MeWo | WM1366 | WM852 | FEMX-V | FEMX-I | WM1382 |
|---|---|---|---|---|---|---|---|---|---|---|---|---|---|---|---|---|---|---|---|
| **RAS/RAF/MEK/ERK** | 1 | 2 | 6 | 3 | 3 | 3 | 8 | 5 | 3 | 5 | 1 | 5 | 2 | 14 | 10 | 1 | 2 | 3 | 2 |
| **WNT** | | | 1 | | | | 1 | 1 | | 1 | | | | 1 | | | | | |
| **hippo** | | 1 | | | | | 1 | | | | | | | 1 | 1 | | | | |
| **PI3K/AKT/mTOR** | 1 | 1 | 1 | | | | 1 | 2 | 1 | 2 | 2 | | 1 | 2 | 1 | 1 | | | |
| **p53/DNA repair** | 2 | 2 | | | 1 | 1 | | 3 | 2 | 2 | 1 | | 1 | 4 | 4 | 1 | 2 | 2 | 1 |
| **nucleosome remodeling** | 1 | | 2 | 1 | | | 1 | | | | | 1 | | 2 | 1 | | 1 | 1 | |

**Supplementary Fig. 12 Oncogenic mutations in human and murine B16-F10 melanoma.** Probable cancer driver mutations in 18 human and murine B16-F10 melanoma cell lines identified upon analysis of RNA and DNA gene panel sequencing data, and after whole-exome sequencing of B16-F10 cells[23] listed with the genes with the most frequent mutations at the top[51]. Stop gained mutations are indicated in red, missense mutations in blue, and splice variant mutation in green. The *MC1R* variants are assumed to be germ-line. The bottom table categorizes the mutations into the most frequently affected signaling pathways for each cell line as indicated below. The $BRAF^{C264R}$ mutation in B16-F10 is not homologous to human the $BRAF^{V600E}$ mutation and is indicated in the figure.

#### Supplementary Fig. 13

### Upstream regulators for YAP$^{high}$ versus hippo$^{low}$ in untreated samples

| upstream regulator | expression log ratio | molecule type | activation z-score | p-value of overlap | target molecules in dataset |
|---|---|---|---|---|---|
| **MITF** | -2.8 | transcription regulator | -7.1 | 1.4E-49 | ACP5,ASAH1,ATP1A1,BEST1,CAPN3,CDK5R1,CHKA,CXCL8,DCT,DSTYK,EDNRB,ESRP1,FRMD4B,GM2A,GNPTAB,GPM6B,GPNMB,GPR137B,GREB1,GZMB,INPP4B,IRF4,ITPKB,IVNS1ABP,KAZN,LGALS3,LYST,MC1R,MICAL1,MITF,MLANA,MMP14,OSTM1,PHACTR1,PIR,PMEL,QDPR,RHOQ,RRAGD,SCARB1,SEMA6A,SHTN1,SLC45A2,SLC7A8,SORT1,SOX13,SOX6,ST3GAL6,STX7,STXBP1,TBC1D16,TMCC2,TMEM251,TNFRSF14,TRPM1,TYR,UBL3,USP48,ZFYVE16 |
| **NFATC2** | -1.7 | transcription regulator | 2.8 | 1.0E-05 | ACP5,CXCL3,GPNMB,INHBA,IRF1,IRF4,MITF,MLANA,NR4A1,OASL,PMEL,STAT5A,TCF4 |
| **SREBF1** | 0.13 | transcription regulator | -2.3 | 2.3E-05 | ACADS,ADGRG1,ATOX1,BEST1,BHLHE41,FDFT1,GPNMB,HSPA1A/HSPA1B,IL1A,LGALS3,SC5D,SCARB1,SLC22A4,STX1A,STXBP1,TRIM63 |
| **TP53** | 0.40 | transcription regulator | 2.0 | 5.0E-04 | ALDH4A1,AMOTL2,AREG,ATP1A1,BDNF,BHLHE41,CLU,CNN2,CTNNB1,CTSF,CXCL8,CYR61,EPHA2,FAT2,FDFT1,FERMT2,FGF2,FSTL3,FUBP1,GDA,GSN,HMGA1,HSPA1A/HSPA1B,IL1A,INHBA,KAT2B,LASP1,LGALS3,MAD1L1,MCL1,MYO10,NCOR2,NDRG2,NKD1,PARD6G,PMAIP1,PODXL,SCPEP1,SEMA6A,SFN,SLC25A13,SMURF2,TCF4,TIMP2,TUBB4A,UBL3,USP48 |
| **TFEB** | -0.01 | transcription regulator | -2.0 | 4.5E-04 | CTSF,HEXA,NAGLU,SCPEP1,TYR |
| **HOXB9** | 1.28 | transcription regulator | 2.0 | 1.8E-04 | AREG,CXCL8,FGF2,NRG1 |
| **NR3C1** | 0.50 | ligand-dependent nuclear receptor | -1.9 | 6.0E-05 | AMIGO2,ATP6AP1,BDNF,CLCF1,CXCL3,CXCL8,CYP27A1,CYR61,DPP7,ERRFI1,IL11,IL1A,IL1RAP,IL7R,INHBA,IRF1,LYST,MAP3K14,MCL1,MMP8,OASL,RBMS2,SCARB1,SFTPC,STAT5A,TNFRSF12A,TRAF5,ZNF704 |
| **RUNX3** | -0.76 | transcription regulator | -1.9 | 5.1E-05 | COL12A1,CRIM1,CYR61,DIP2C,EXT1,FGF2,FZD2,NUAK1,SGCD,TPM2 |
| **SNAI1** | -0.80 | transcription regulator | 1.2 | 6.8E-06 | AXL,CXCL8,CYR61,ESRP1,FLNB,GSN,LASP1,MMP14,PARVB,PEBP1,TPM1 |
| **CTNNB1** | -0.72 | transcription regulator | -1.2 | 2.0E-05 | ADRA2C,ALDH1A1,APOD,CLU,CNN2,CTNNB1,CTSF,CXCL8,CYR61,DCT,FSTL3,GJB1,GPR137B,HMG20B,ID4,IRF4,LAMC2,LGALS3,MCL1,MITF,MMP14,NDRG2,NKD1,NR4A1,NRG1,OSBPL1A,PKP3,SFN,STAT5A,STXBP1,TCF4,TNIK |
| **MEOX2** | - | transcription regulator | -1.1 | 6.9E-06 | CXCL3,CXCL8,FGF2,ID4,MMP14,PTX3,VEGFC |
| **SOX10** | -4.2 | transcription regulator | -1.1 | 2.1E-04 | EDNRB,GDA,GZMB,MITF,PLP1 |
| **WT1** | 1.3 | transcription regulator | 1.0 | 4.8E-04 | AREG,CTNNB1,FDFT1,GSN,IL11,IL1RAP,LGALS3,MCL1,MMP14,PIR,PODXL,SMAD3,ZNF280B |
| **SOX2** | -0.80 | transcription regulator | -1.0 | 3.8E-04 | ABCC3,ALDH1A1,CITED1,CTNNB1,EPHA2,GAB2,GATA4,GREB1,INHBA,PARD6G,SALL4,SOX13,SOX6,TEAD3,TXNRD1,VEGFC |
| **NFKBIA** | 0.68 | transcription regulator | 0.9 | 6.7E-07 | AXL,CLU,CTNNB1,CTSF,CXCL3,CXCL8,FGF2,GM2A,GPR176,GZMB,HMGA1,IGFBP6,IL11,IL1A,IL7R,IRF1,MMP14,NAGLU,NDRG2,NR4A1,PTX3,SORL1,SPTBN1,TIMP2,VEGFC |
| **MYCN** | - | transcription regulator | -0.9 | 8.1E-04 | ABCC3,ABCD1,ALDH1A1,AMOTL2,BASP1,CLU,CRIM1,GATA4,HMGA1,INHBA,MXI1,PMAIP1,TIMP2,TPM1 |
| **SP1** | 0.17 | transcription regulator | -0.9 | 3.1E-04 | BACE2,BDNF,CXCL3,CXCL8,EDA,FGF2,HMGA1,ID4,IL1A,IRF1,IRF4,MCL1,MECP2,MMP14,PDGFC,PMAIP1,ROBO4,SCARB1,SLC22A4,SMAD3,STX1A,TIMP2,TXNRD1 |
| **JUN** | 1.0 | transcription regulator | 0.8 | 3.2E-04 | ACP5,AXL,BDNF,CDK5R1,CHL1,CLU,CXCL3,CXCL8,CYR61,FGF2,GREB1,HMGA1,IGFBP6,IL1A,IL7R,LGALS3,MMP8,MXI1,NR4A1,PTX3 |
| **TP63** | - | transcription regulator | -0.8 | 8.8E-04 | AREG,AXL,CTNNB1,CXCL8,CYR61,EPHA2,FUBP1,IGFBP6,IL1RAP,INHBA,MLPH,PMAIP1,SFN,SMURF2,STX1A,TPM1,USP48 |
| **ESR1** | - | ligand-dependent nuclear receptor | -0.6 | 7.6E-04 | ABCC3,AREG,ARSG,CDK5R1,CERS1,COL13A1,CRIM1,CTNNB1,CXCL3,CXCL8,EFNB2,FARP2,FZD2,GJB1,GM2A,GREB1,GSN,IFI44L,IL1A,IRF1,IRF4,MAP3K14,MC1R,PMAIP1,PTX3,RENBP,RHOQ,SCARB1,SCUBE2,SLC1A4,SLC6A8,SLC7A8,SMAD3,SMURF2,SPTBN1,STAT5A,TIMP2,TPM1,TRAK1,UBL3,VEGFC |
| **STAT6** | 0.16 | transcription regulator | -0.5 | 7.0E-04 | ACP5,AREG,DDAH1,EXT1,FNIP2,GPM6B,IL1A,IRF1,IRF4,LDLRAD3,MMP14,PDGFC,SOX13,TNIK |
| **ATF4** | 0.18 | transcription regulator | 0.5 | 4.7E-04 | AREG,CTNNB1,CYP27A1,LGALS3,MCL1,PMAIP1,PTX3,SLC1A4,SLC6A9,TNFRSF12A |

**Supplementary Fig. 13 IPA core analysis of RNA sequencing data identifies MITF as the top upstream regulator separating the baseline YAP$^{high}$ versus the YAP$^{low}$ groups.** Differentially expressed genes, when comparing YAP$^{high}$ versus YAP$^{low}$ cell lines (see Fig. 4b), with a corrected *P* value of <0.05 were used in an IPA analysis for identifying upstream regulator components. The table is displaying the identified upstream regulators with an activation z-score >0.5 or <-0.5 and *P* value of overlap <0.0005. Expression log ratio: Log2-value YAP$^{high}$ versus YAP$^{low}$ TPMs for transcription of the indicated upstream regulator. A low or high value indicates transcriptional difference of the upstream regulator itself. Molecule type: Depicts the biological function of the upstream regulator. Target molecule in dataset: Lists all differently expressed genes in the dataset that are linked to the upstream regulator. MITF was identified as the top upstream regulator separating the baseline YAP$^{high}$ versus the YAP$^{low}$ groups.

#### Supplementary Fig. 14

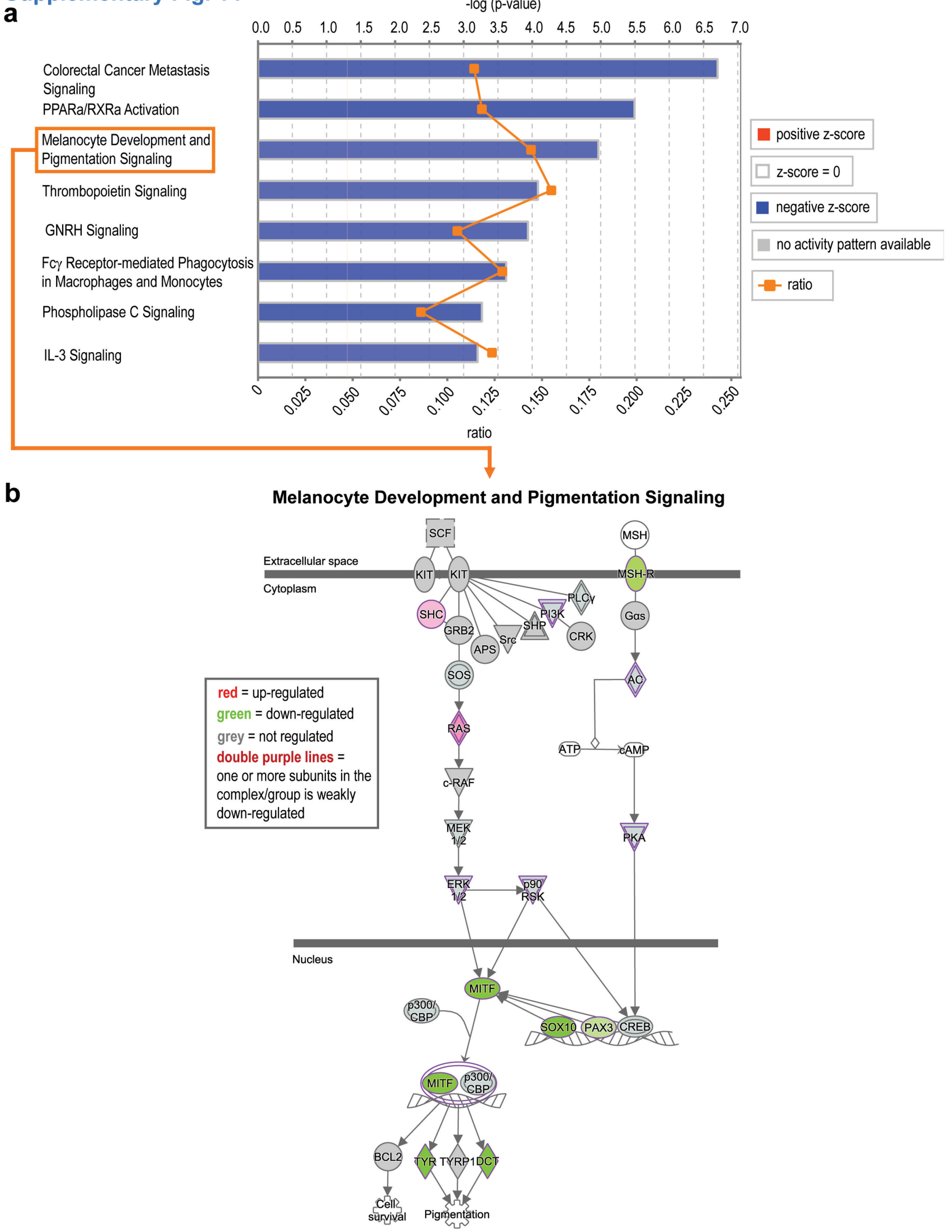

**Supplementary Fig. 14 IPA core analysis of RNA sequencing data identifies canonical pathways separating the baseline YAP$^{high}$ versus the YAP$^{low}$ groups. a**, Differentially expressed genes, when comparing YAP$^{high}$ versus YAP$^{low}$ cell lines (see Fig. 4b), with a corrected *P* value of <0.05 were analyzed for direct relationships in a IPA core analysis. The plot is displaying the eight top canonical pathways identified with core analysis with a -log(*P* value) >3 and an absolute z-score >2. Negative z-score is shown as blue bars along the -log(*P* value) axis. The ratio of genes in this dataset matching the pathway, divided by the total genes in the pathway, is shown as the orange line. **b**, Pathway map for the IPA analysis-identified "Melanocyte Development and Pigmentation Signaling pathway". Up-regulated RNA expression is indicated in red, down-regulated in green, no regulation in grey and double purple lines indicate that transcription of one or more of the components of the depicted protein complex is down-regulated.

#### Supplementary Fig. 15

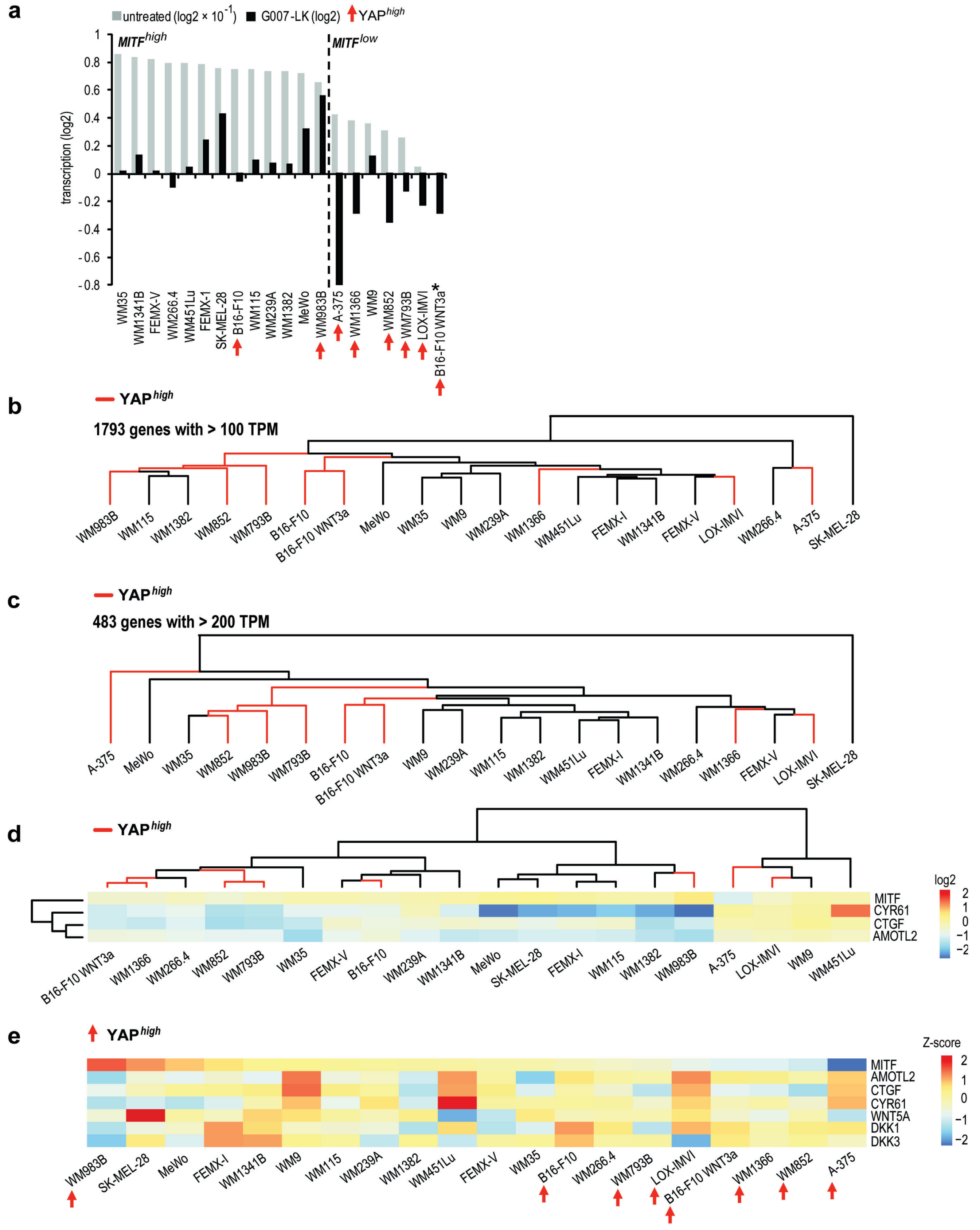

**Supplementary Fig. 15 High relative baseline YAP signaling activity correlates with low *MITF* expression and G007-LK-induced changes in *MITF* expression cannot be explained alone by changes in YAP or WNT signalling activity. a**, Bar chart showing *MITF* expression in untreated samples (grey bars sorted descending from left to right, log2-transformed TPMs × $10^{-1}$) and upon treatment with G007-LK (1 µM) for 24 hours (black bars, log2 values from treated versus untreated TPMs)(see also Fig. 4b). * indicates that no untreated value is inserted for this sample. The dotted vertical line depicts division between samples with increased ($MITF^{high}$, left) or decreased ($MITF^{low}$, right) expression of *MITF* upon tankyrase inhibitor treatment. For **a**-**e**: Change in gene expression for 18 G007-LK-treated (1 µM) human and murine B16-F10 melanoma cell lines is shown. Samples displaying high relative transcription of YAP signaling target genes ($YAP^{high}$, see Fig. 4a) are highlighted by orange arrows or by orange branches in the dendrogram. B16-F10 WNT3a = WNT3a + G007-LK relative to WNT3a-stimulated control. **b**, Clustering of 1793 genes with changed gene expression with >100 TPMs in >80% of the comparisons (log2). **c**, Clustering of 483 transcribed genes with >200 TPMs (log2). **d**, Heatmap and clustering of transcribed YAP signaling target genes (*Cyr61*, *Ctgf* and *Amotl2*) versus *MITF* upon G007-LK treatment. Decreased YAP signaling ($YAP^{decreased}$) does not correlate decreased *MITF* transcription ($MITF^{decreased}$). Scale bar indicates relative differences in log2 TPMs. **e**, Heatmap and clustering showing that altered levels of YAP signaling activity (*Cyr61*, *Ctgf* and *Amotl2*) or expression of DKK3, WNT5a and DKK1 do not coincide with decreased *MITF* transcription (sorted descending from right to left).

**Supplementary Fig. 16**

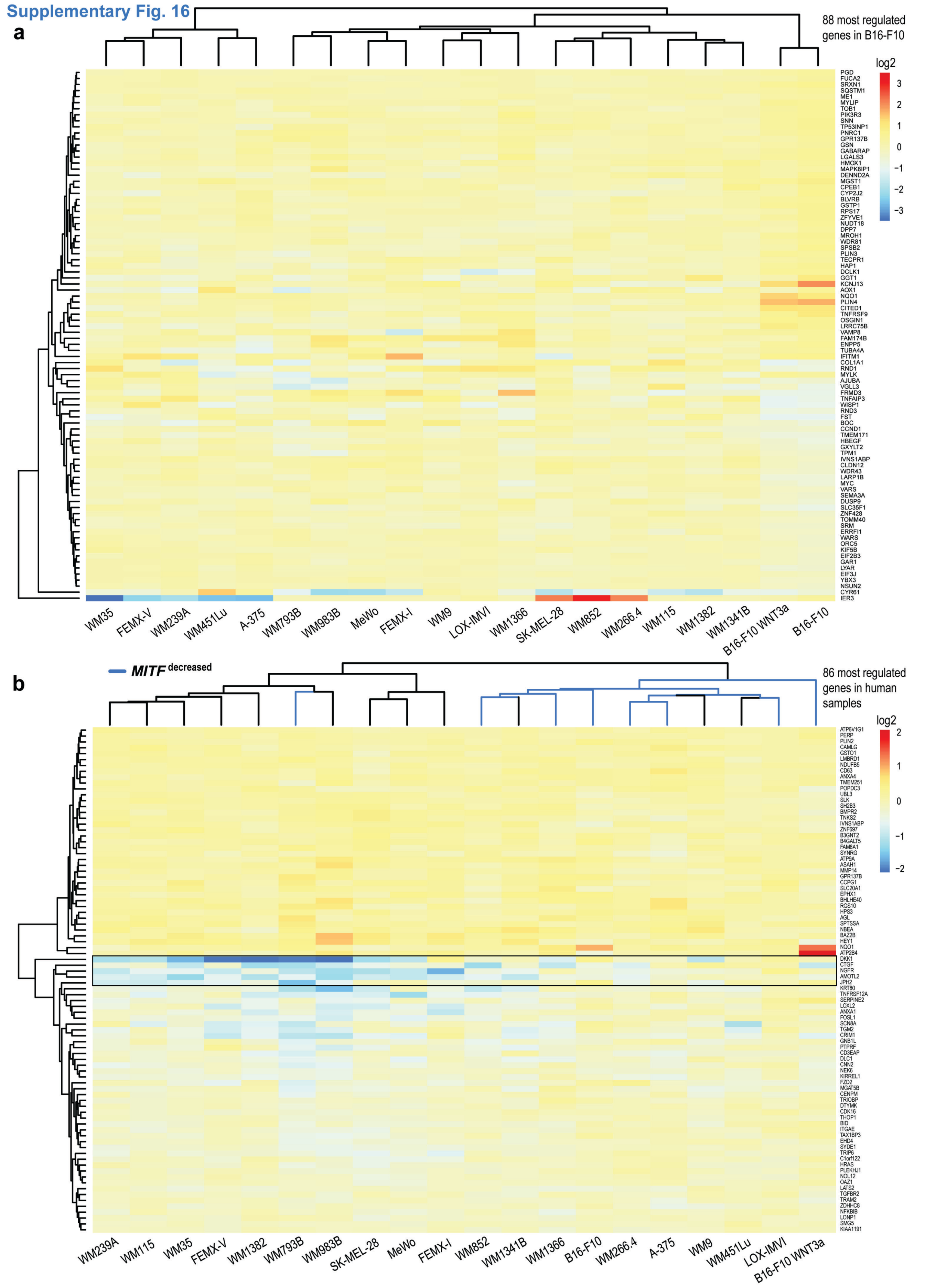

**Supplementary Fig. 16 Top lists for G007-LK-induced regulation of gene expression.** **a**, The 88 most regulated genes in B16-F10 samples. Murine B16-F10 samples do not cluster with human samples. For **a** and **b**: Heatmap and clustering of change in gene expression for 18 G007-LK-treated (1 µM) human and murine B16-F10 melanoma cell lines. Scale bar indicates log2-values. B16-F10 WNT3a = WNT3a + G007-LK relative to WNT3a-stimulated control. **b**, The 86 most regulated genes in human samples. The clustering is in particular orchestrated by changes in expression of *DKK1*, *CTGF*, *NGFR*, *AMOTL2* and *JPH2* (highlighted by black box). 7 of 8 samples in the $MITF^{decreased}$ subset (see Fig. 4b), including the B16-F10 samples, are found on the right side of the heatmap (highlighted by blue branches in the dendrogram).

#### Supplementary Fig. 17

**a**

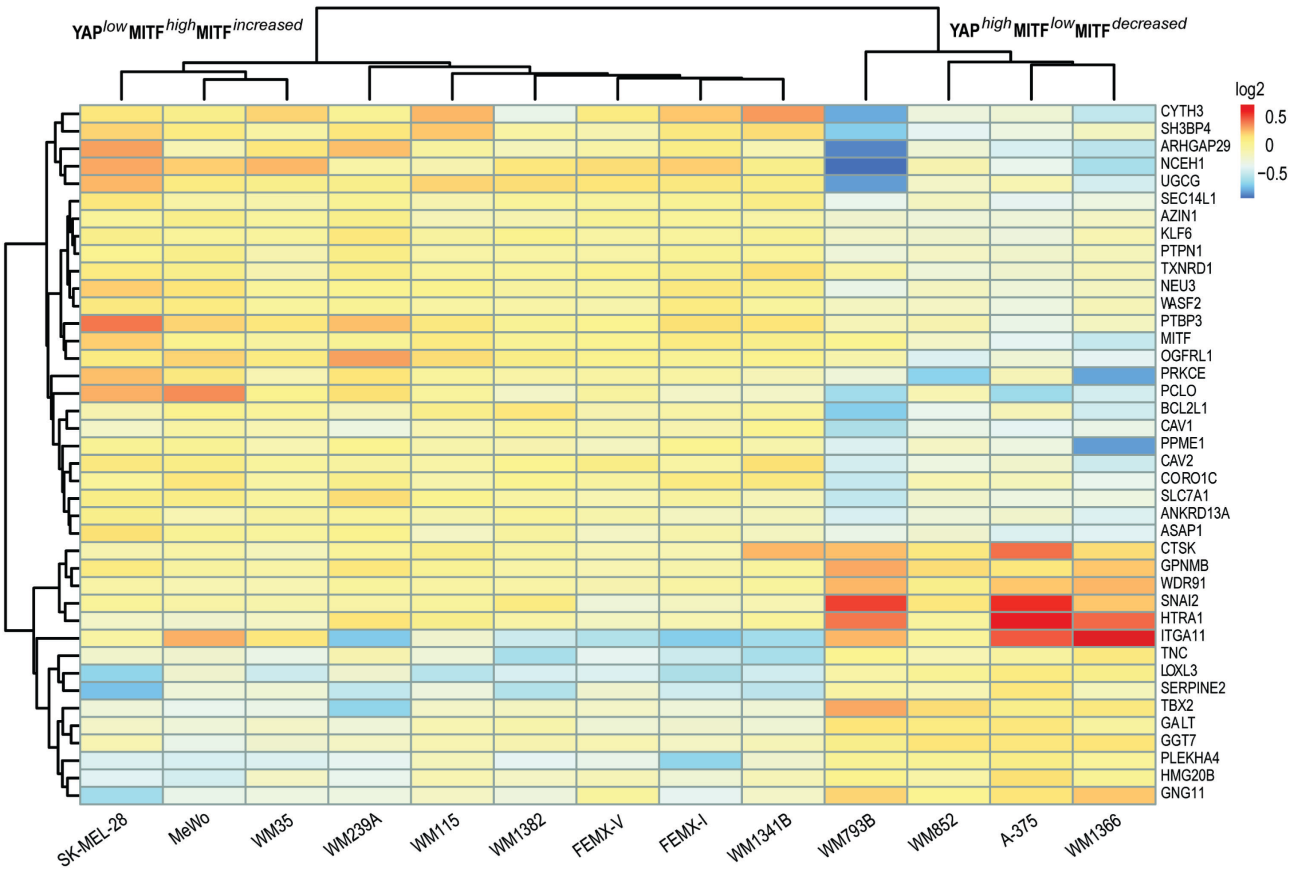


**b**

| upstream regulator | expression log ratio | molecule type | activation z-score | p-value of overlap | target molecules in dataset |
|---|---|---|---|---|---|
| **TCF7L2** | 0.061 | transcription regulator | -2.6 | 6.6E-03 | ANKRD13A,DOCK9,EPAS1,ERBIN,GCLM,NPC1,SNAI2,STK39 |
| **KLF3** | -0.12 | transcription regulator | 2.3 | 2.0E-03 | DYRK3,EXTL1,HLCS,HPS1,MSN,OGFRL1,TES,TMOD3,UGCG |
| **PGR** | 1.46 | ligand-dependent nuclear receptor | -1.5 | 5.0E-04 | BCL2L1,CD36,DDX21,EPAS1,GPRC5A,NPC1,SNAPC1,TNC,UGCG |
| **AR** | -0.17 | ligand-dependent nuclear receptor | -1.4 | 2.4E-03 | BCL2L1,CAV1,CAV2,DSG2,EPAS1,MEF2C,NPC1,PROS1,SNAI2,STK39,TRIM25 |
| **GLI1** | 0.41 | transcription regulator | -1.3 | 1.5E-03 | BCL2L1,CAV1,MEF2C,NR2F2,PRKCE,PVR,TNC |
| **ESR1** | 0.69 | ligand-dependent nuclear receptor | -1.1 | 3.7E-07 | BCL2L1,CAV1,CAV2,DAB2,DDB2,DDX21,DLG1,DSG2,EEA1,EXOC5,FOSL2,GNAQ,GNG12,HPS1,HTRA1,MAP3K20,NQO1,PDLIM5,PROS1,PVR,RGS3,SEC14L1,SLC7A1,SNAI2,SSR3,TACC1,TMOD3,TNFAIP3,TRIO,WASF2 |
| **NFE2L2** | 0.051 | transcription regulator | -1.0 | 6.7E-04 | BCL2L1,CD36,CTSK,EPAS1,GCLM,MEF2C,NFE2L1,NQO1,PTPN1,SNAI2,TXNRD1 |
| **ERG** | -2.4 | transcription regulator | -1.0 | 1.4E-03 | CLIP1,ETS1,MAP3K5,NR4A2,RGS3,TACC1,WDR91 |

**Supplementary Fig. 17 Tankyrase inhibition induces a transcriptional program subdividing the YAP$^{low}$*MITF*$^{high}$*MITF*$^{increased}$ and YAP$^{high}$*MITF*$^{low}$*MITF*$^{decreased}$ groups. a**, Change in gene expression for G007-LK-treated (1 µM) YAP$^{low}$*MITF*$^{high}$*MITF*$^{increased}$ (left) versus YAP$^{high}$*MITF*$^{low}$*MITF*$^{decreased}$ (right) subgroups of human cell lines (see Fig. 4e). Differentially expressed genes with adjusted *P* value <0.01 are shown. Scale bar indicates relative differences in log2 counts. **b**, IPA core analysis identifies *TCF7L2* (based on z-score) and *ESR1* (based on *P* value of overlap) as the top upstream regulator separating the in YAP$^{low}$*MITF*$^{high}$*MITF*$^{increased}$ and YAP$^{high}$*MITF*$^{low}$*MITF*$^{decreased}$ subgroups. Differentially expressed genes with an adjusted *P* value of <0.1 were used in an analysis for identifying upstream regulator components. The table is displaying the identified upstream regulators with an activation z-score >1 or <-1 and *P* value of overlap <0.01. Expression log ratio: Log2 values (counts) for YAP$^{low}$*MITF*$^{high}$*MITF*$^{increased}$ versus YAP$^{high}$*MITF*$^{low}$*MITF*$^{decreased}$ indicating transcription of the indicated upstream regulator. A low or high value indicates transcriptional difference of the upstream regulator itself. Molecule type: Depicts the biological function of the upstream regulator. Target molecule in dataset: Lists all differently expressed genes in the dataset that are linked to the upstream regulator.

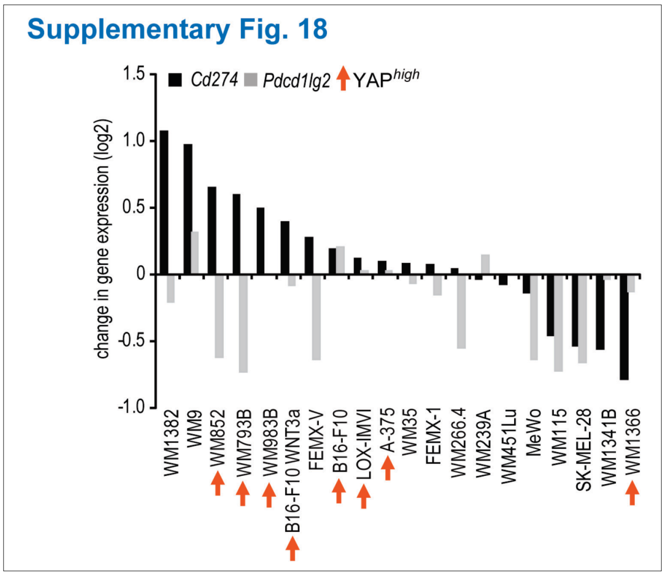


**Supplementary Fig. 18 G007-LK treatment can regulate *Cd274 and Pdcd1lg2* expression in melanoma cell lines.** Bar chart showing *Cd274* (*PD-L1*, black bars sorted descending from left to right) and *Pdcd1lg2* (*PD-L2, grey bars*) expression (log2) in samples upon treatment with G007-LK (1 μM) for 24 hours. B16-F10 WNT3a = WNT3a + G007-LK relative to WNT3a-stimulated control. Samples displaying high relative baseline transcription of YAP signaling target genes (YAP$^{high}$, see Fig. 4a) are highlighted by orange arrows.

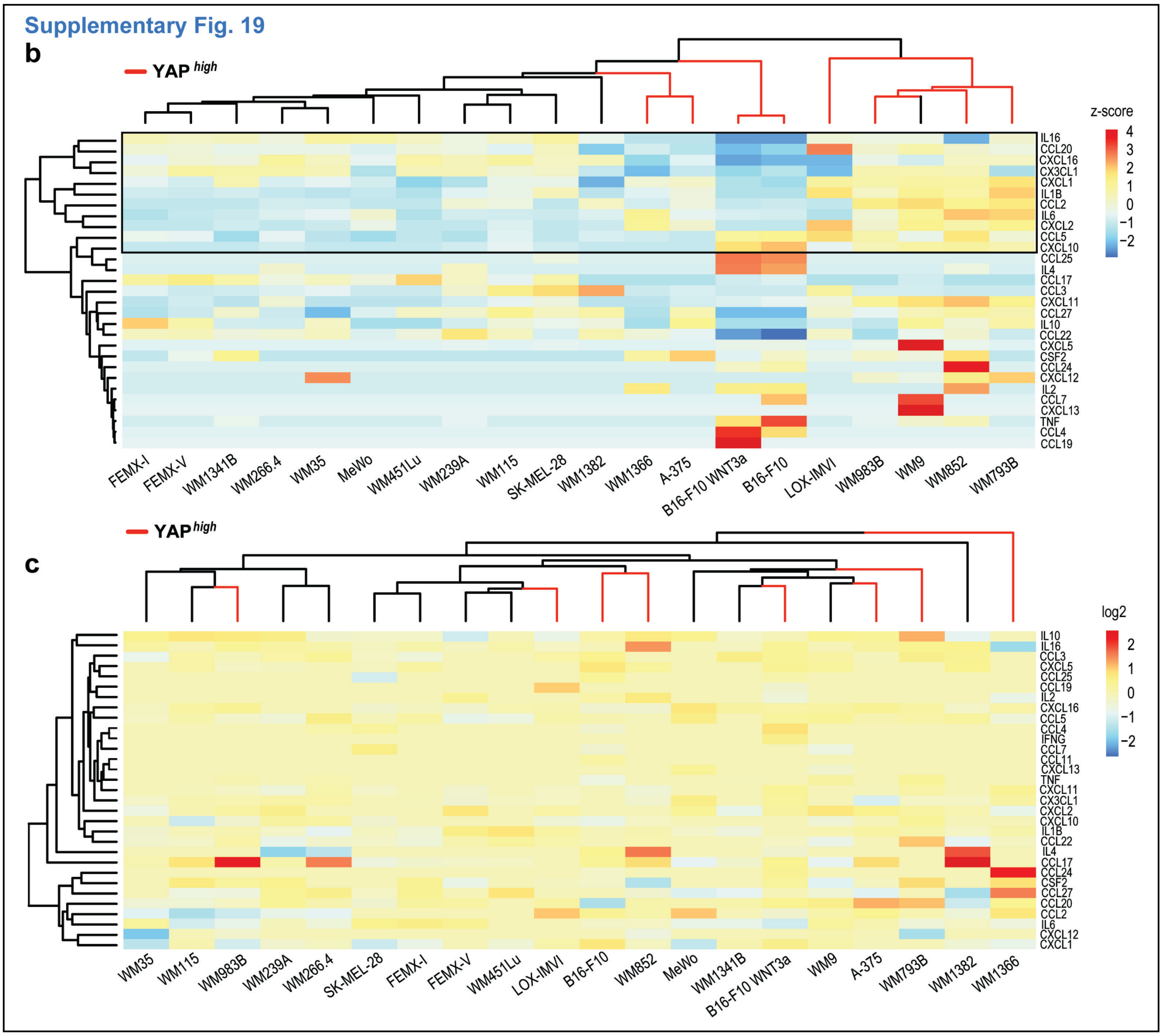


**Supplementary Fig. 19 YAP signaling activity correlates with baseline cytokine and chemokine expression *in vitro* and is moderately regulated by G007-LK treatment. a**, Heatmap and clustering of transcribed cytokines and chemokines for 18 human and murine B16-F10 melanoma cell lines. YAP$^{high}$ samples cluster partly and the clustering is in particular orchestrated by different baseline expression of *IL16, CCL20, CXCL16, CX3CL1, CXCL1, IL1B, CCL2, IL6, CXCL2, CCL5 and CXCL10* (highlighted by black box). Scale bar indicates z-score values. For **a** and **b**: B16-F10 WNT3a = WNT3a + G007-LK relative to WNT3a-stimulated control. Samples displaying high relative baseline transcription of YAP signaling target genes (YAP$^{high}$, see Fig. 4a) are highlighted by orange branches in the dendrogram. List of analyzed cytokines and chemokines are taken from Bio-Plex Pro Mouse Chemokine Panel 33-plex (12002231, Bio-Rad). **b**, Heatmap and clustering of change in gene expression for 18 G007-LK-treated (1 µM) human and murine B16-F10 melanoma cell lines. Scale bar indicates log2-values. Moderate regulation of cytokine and chemokine transcription was observed upon G007-LK treatment in the *in vitro* cultivated cell lines.
